# Hydrophobic tails enable diverse functions of the extracellular chaperone clusterin

**DOI:** 10.1101/2024.10.30.620894

**Authors:** Patricia Yuste-Checa, Alonso I. Carvajal, Chenchen Mi, Sarah Paatz, F. Ulrich Hartl, Andreas Bracher

## Abstract

Clusterin, a conserved secretory glycoprotein abundant in blood plasma and cerebrospinal fluid, functions as a molecular chaperone and apolipoprotein (Wyatt et al. 2013, Raulin et al. 2022). Dysregulation of clusterin is linked to late-onset Alzheimer’s disease, cardiovascular pathology and cancer (Rohne et al. 2016, Satapathy and Wilson 2021, Wilson et al. 2023). Despite its prominent role in extracellular proteostasis, the chaperone mechanism of clusterin has remained unclear. Here we present crystal structures of human clusterin, revealing a discontinuous three-domain architecture. Structure-based mutational analysis demonstrated that two intrinsically disordered, hydrophobic peptide tails enable diverse clusterin activities. Resembling the N-terminal substrate binding regions of so-called small heat shock proteins, these sequences mediate clusterin’s chaperone function in suppressing amyloid-β, tau and α-synuclein aggregation. In conjunction with highly conserved surface areas, the tail segments also participate in clusterin binding to very low density lipoprotein receptor (VLDLR) and cellular uptake. Moreover, the disordered tails cooperate with an adjacent amphipathic helix in lipoprotein formation, but remain accessible for chaperone function in the lipoprotein complex. The remarkable versatility of these sequences allows clusterin to function alone or bound to lipid in maintaining solubility of aberrant extracellular proteins and facilitating their clearance by endocytosis and lysosomal degradation.

## Main

Extracellular molecular chaperones are implicated in numerous pathologies including Alzheimer’s disease (AD) and other neurodegenerative disorders associated with aggregate deposition (Wilson, Satapathy and Vendruscolo 2023). The conserved vertebrate secretory glycoprotein clusterin (Clu, also known as Apolipoprotein-J, ApoJ), functions as an abundant extracellular chaperone in mammalian blood plasma, seminal and cerebrospinal fluid (Wyatt, Yerbury et al. 2013, Rohne, Prochnow and Koch-Brandt 2016, Wilson and Zoubeidi 2017, Satapathy and Wilson 2021). Initially described as a factor mediating cell clustering *in vitro* (Blaschuk et al. 1983, Fritz et al. 1983), Clu was also shown to have an immune-regulatory function by inhibiting complement mediated cell lysis (Jenne and Tschopp 1989, Menny et al. 2021) and to fractionate with high-density lipoprotein (HDL)-like particles in plasma and the central nervous system (de Silva et al. 1990, Jenne et al. 1991, Suzuki et al. 2002, Lu et al. 2022, Raulin, Martens and Bu 2022). Characterization of Clu as a molecular chaperone is based on its activity *in vitro* to inhibit the aggregation of non-native proteins (Humphreys et al. 1999) and neurodegenerative disease proteins amyloid-β (Aβ) (Hammad et al. 1997, Yerbury et al. 2007, Narayan et al. 2011, Scheidt et al. 2019, Kim et al. 2022), tau (Wojtas et al. 2020, Yuste-Checa et al. 2021), α-synuclein (Whiten et al. 2018, Yuste-Checa, Trinkaus et al. 2021) and prion protein (McHattie and Edington 1999, Xu et al. 2008). Consistent with a role as chaperone, Clu is upregulated in AD and colocalizes with Aβ deposits in patient brain, possibly reflecting a role in clearance of misfolded proteins by facilitating receptor mediated endocytosis and lysosomal degradation (Wyatt et al. 2011, Yeh et al. 2016, Itakura et al. 2020). Indeed, certain *CLU* gene alleles rank among the most significant genetic risk factors for late onset AD (Harold et al. 2009, Lambert et al. 2009, Jansen et al. 2019).

Clu is synthesized as a ∼50 kDa precursor protein with a secretory signal peptide that is removed upon translocation into the endoplasmic reticulum (ER), where six N-glycans are attached. The protein chain is cleaved into β- and α-chains (residues 23–227 and 228–449, respectively) by furin-like proteases in the Golgi (Supplementary Figure 1a). The chains remain covalently linked by five conserved disulfide bonds (Choi-Miura et al. 1992). To provide a basis for detailed mechanistic studies, here we determined the crystal structure of human Clu and explored its function by rational mutagenesis. We identified two hydrophobic flexible tails as being critical for binding non-native protein, receptor mediated endocytosis and – in conjunction with an amphipathic helix – formation of lipoprotein particles. Interestingly, Clu maintains functionality as chaperone when bound to lipid.

### Crystal structure of clusterin

Human Clu was over-expressed in HEK293E cells and purified from the cell culture supernatant. The protein was monomeric at pH 5 (apparent size ∼90 kDa), but additionally formed dimers of ∼220 kDa at pH 6.5-8.5 (Supplementary Figure 1b) without measurably affecting Clu secondary structure content and stability (Supplementary Figure 1c, d). Purified Clu exhibited N-glycan heterogeneity and almost complete processing into α- and β-chains migrating at ∼40 kDa on SDS-PAGE (Supplementary Figure 1e). For crystallization, we designed a single-chain construct (Clu-Δ(214–238)) comprising a deletion of a predicted unstructured sequence including the furin cleavage site. Upon expression in the presence of the α-mannosidase-I inhibitor kifunensine (Chang et al. 2007) to generate uniform oligo-mannose N-glycans (Supplementary Figure 1e, f), purified Clu-Δ(214–238) crystallized at acidic pH in two distinct lattice types, crystal forms I and II, diffracting to 2.8 and 3.5 Å resolution, respectively (Supplementary Table 1). The structures were solved by molecular replacement using the Alphafold2 model of Clu as a search template (https://www.alphafold.ebi.ac.uk/entry/P10909). Crystal form I contained one copy of Clu-Δ(214–238), crystal form II two independent copies (Figure 1a and Supplementary Figure 2a).

**Figure 1:**
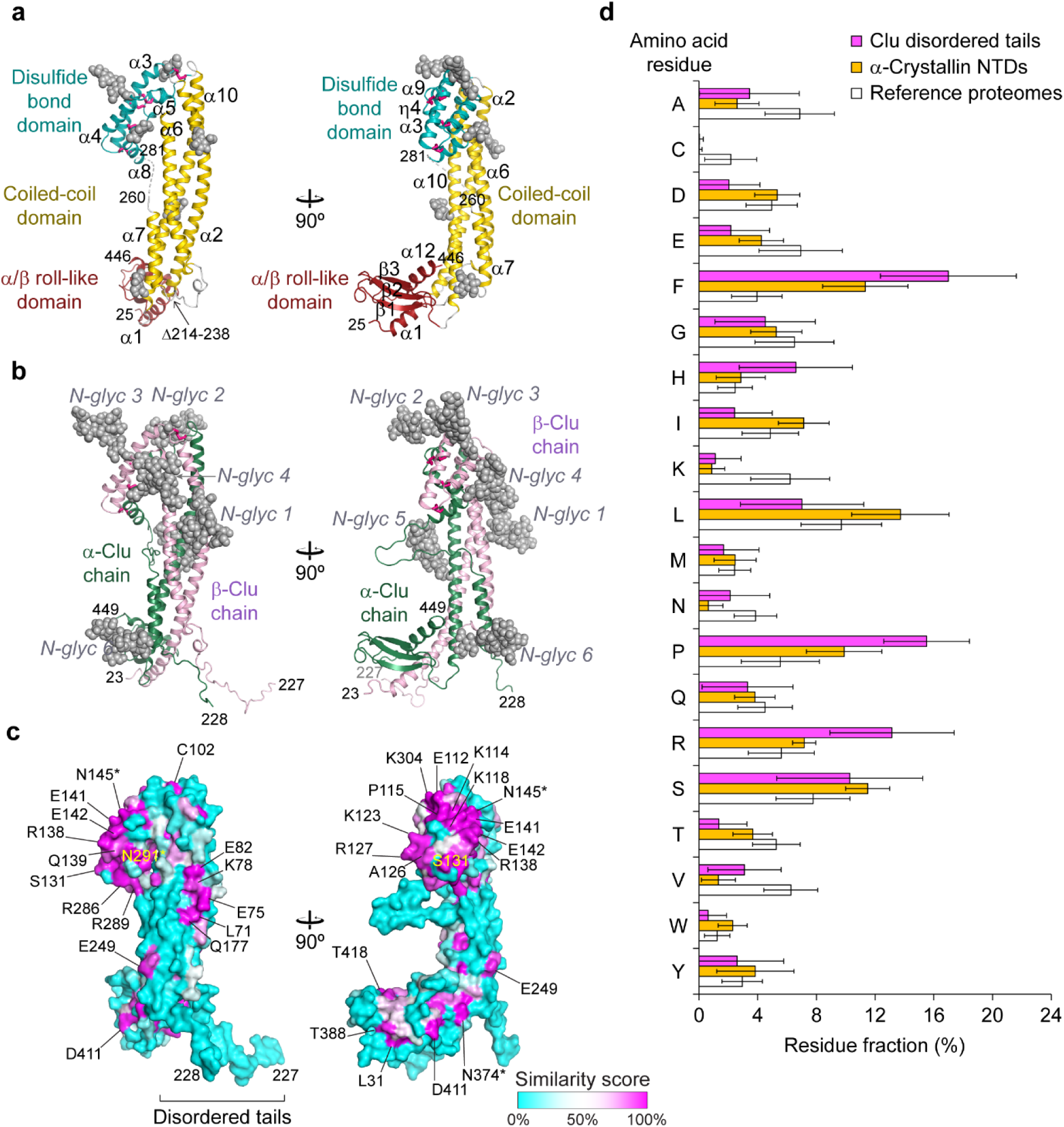
Structure of clusterin. (**a**) Crystal structure of Clu-Δ(214–238). The asymmetric unit of crystal form I is shown in perpendicular views. Coiled-coil, disulfide domain and α/β roll-like domains are indicated in gold, teal and dark red, respectively. The dashed line indicates a disordered segment, residues 261–280. Residue numbering refers to the human Clu precursor sequence. Secondary structure elements are shown in ribbon representation. Disulfide bonds are shown as purple sticks, partially ordered N-glycan structures present in the crystal in space-filling mode in gray. First and last structured residue, the deletion site and secondary structure elements of Clu are indicated. (**b**) Structural model for wild-type Clu. The Clu β- and α-chains are colored pink and dark green, respectively. The unstructured residues 204–227, 228–238 and 261–280 have been modeled in arbitrary conformations. The full N-glycans (N-glyc 1-6, gray spheres) have been modeled as oligo-mannose trees, i. e. before diversification by N-glycan processing in the Golgi. (**c**) Sequence conservation mapped onto the molecular surface of Clu. The same views as in Figure 1b are shown. A cyan-white-magenta color gradient indicates increasing surface conservation based on the similarity score from the sequence alignment shown in Supplementary Figure 4. Highly conserved residues are indicated. Asterisks indicate highly conserved N-glycosylation sites. (**d**) Sequence bias in the unstructured tails of Clu. A histogram of the amino acid composition of residues 204–238 (human Clu numbering) in 242 representative Clu sequences is shown in magenta. For comparison, the average compositions of the N-terminal domains (NTDs) in α-crystallin A and B homologs (orange; 662 CRYAA/HSPB4 and 444 CRYAB/HSPB5 sequences) and the average composition of selected vertebrate reference proteomes (white; human, opossum, platypus, chicken, clawed frog, coelacanth, zebrafish and ghost shark) are shown. Error bars designate standard deviations.

The three conformations in the crystal lattices differed slightly in their domain orientations, with r.m.s.d. values of 1.5-2.0 Å (C-α positions) (Supplementary Figure 2b). The Alphafold2 model showed larger deviations, mainly due to domain reorientations, with r.m.s.d. values of 4.7-5.6 Å (C-α positions) (Supplementary Figure 2c).

The Clu structure is composed of three domains, a coiled-coil, a small helical domain containing all disulfide bridges – in the following referred to as disulfide domain (DD) – and a C-terminal α/β roll-like domain (Figure 1a). The ∼85 Å long coiled-coil bundle is composed of helices α2 and α6 from the β-chain and α10 from the α-chain in wild-type (WT) Clu. The coiled-coil interactions of these helices form the majority of inter-chain contacts (Figure 1a). Helix α2 runs anti-parallel to helices α6 and α10. The shorter 7-turn helix α7 aligns with the bundle, widening the distance between helices α6 and α10. The five inter-chain disulfides connect helix α3 and its flanking linkers with the short helices α8, η3 and α9, which zig-zag along helix α3 (Figure 1a). Helices α4 and α5 buttress these helices in the α-chain. While the DD seems fairly rigid, its linkers to helices α2 and α10 deviate considerably between crystal forms, indicating structural plasticity (Supplementary Figure 2b). The α/β roll-like domain consists of a three-stranded anti-parallel β-sheet, which wraps around helix α12 (Figure 1a). Sheet and helix are flanked by helix α1. The α/β roll-like domain thus also contributes to the inter-chain contacts in WT-Clu.

The N-glycans in Clu-Δ(214–238) point towards solvent channels in the crystal lattices and are partially ordered (grey spheres, Figure 1a). These hydrophilic elements function presumably in stabilizing the protein in solution. Three of the N-glycans are located in the DD, while the other three are arranged along the coiled-coil bundle, together making up ∼30% of protein mass (Figure 1b and Supplementary Figure 1e). Importantly, the model of the WT structure contains extended unstructured regions, including disordered tails at the C-terminus of the β-chain (residues 199–227) and at the N-terminus of the α-chain (residues 228–244). These tails are generated by furin cleavage in the unstructured sequence of WT-Clu that is deleted in Clu-Δ(214–238). The α7-α8 loop (residues 261–280) is also disordered (Figure 1a). Thus, helix α7 (residues 244–257) is flanked by flexible regions in WT-Clu.

To rule out structural artifacts resulting from the deletion in the crystallized construct, we analyzed both Clu-Δ(214–238) and WT-Clu by hydrogen-deuterium exchange combined with mass spectrometry (H/DX-MS) at pH 5, where the proteins are monomeric (Supplementary Figure 1b, f and 3). WT-Clu generally displayed slow deuterium incorporation in regions that formed secondary structure in Clu-Δ(214–238) (Supplementary Figure 3a-c). Residues 24–33 at the N-terminus, residues 191–195, 220–226 and 234–240 in the predicted flexible tails, as well as residues 270–276 in the disordered α7-α8 loop showed rapid deuterium incorporation in both WT-Clu and Clu-Δ(214–238) (Supplementary Figure 3b,c), consistent with the structural model of WT-Clu derived from the Clu-Δ(214–238) crystal structures (Figure 1b).

### Regions of functional relevance

Clu homologs share ∼15% sequence identity and ∼40% similarity (Supplementary Figure 4). Areas of high surface conservation are restricted to the DD (Figure 1c). We noticed that the disordered tails at the C-terminus of the β-chain (β-tail) and the N-terminus of the α-chain (α-tail) (Figure 1b), while lacking sequence conservation, display a clear compositional bias towards the hydrophobic amino acid Phe (Figure 1c, d). Other residues over-represented in this region are His, Arg, Pro and Ser, with Pro and Ser known to be generally enriched in loop regions (Liu et al. 2002). Lys and Cys residues were less frequent than in reference proteomes. Interestingly, a similar sequence bias is found in the flexible N-terminal domains (NTDs) of small heat shock protein (sHsp) chaperones, such as α-crystallin from vertebrates, which are known to interact with non-native client proteins (Jaya et al. 2009, Kriehuber et al. 2010, Woods et al. 2023) (orange bars in Figure 1d). Hydrophobic clefts reminiscent of substrate protein binding sites in molecular chaperones such as Hsp60 and Hsp70 are absent in the Clu crystal structure. However, we detected two areas of hydrophobic contacts between Clu molecules in the crystal lattice (Supplementary Figure 5a, b): Val26, Met34, Gln37 and Val389 in the α/β roll-like domain flank a hydrophobic crystal contact, while Ile46, Val50, Val369, Ala373 and Thr376 in the coiled-coil domain contact the artificial peptide linkage in Clu-Δ(214–238). Moreover, the narrow cleft between the DD and the coiled-coil might provide a binding site for an extended peptide after slight opening.

In addition to Clu-Δ(214–238), henceforth referred to as tail mutant TL1, we generated a series of Clu mutant proteins targeting the flexible tail sequences (mutants TL2-TL4), conserved surface residues in the disulfide domain (DD1-DD3), the amphipathic helix α7 (CC1) and putative substrate binding regions in the coiled-coil domain (CC2-CC4) and the α/β roll-like domain (AB1, AB2) (Figure 2a). In case of DD1 and DD2 we introduced multiple substitutions to maximize functional effects, while maintaining overall surface properties such as net charge and hydrophobicity. CD spectra indicated helical structure and thermal stability similar to WT-Clu for most mutants (Supplementary Figure 5c, d).

**Figure 2:**
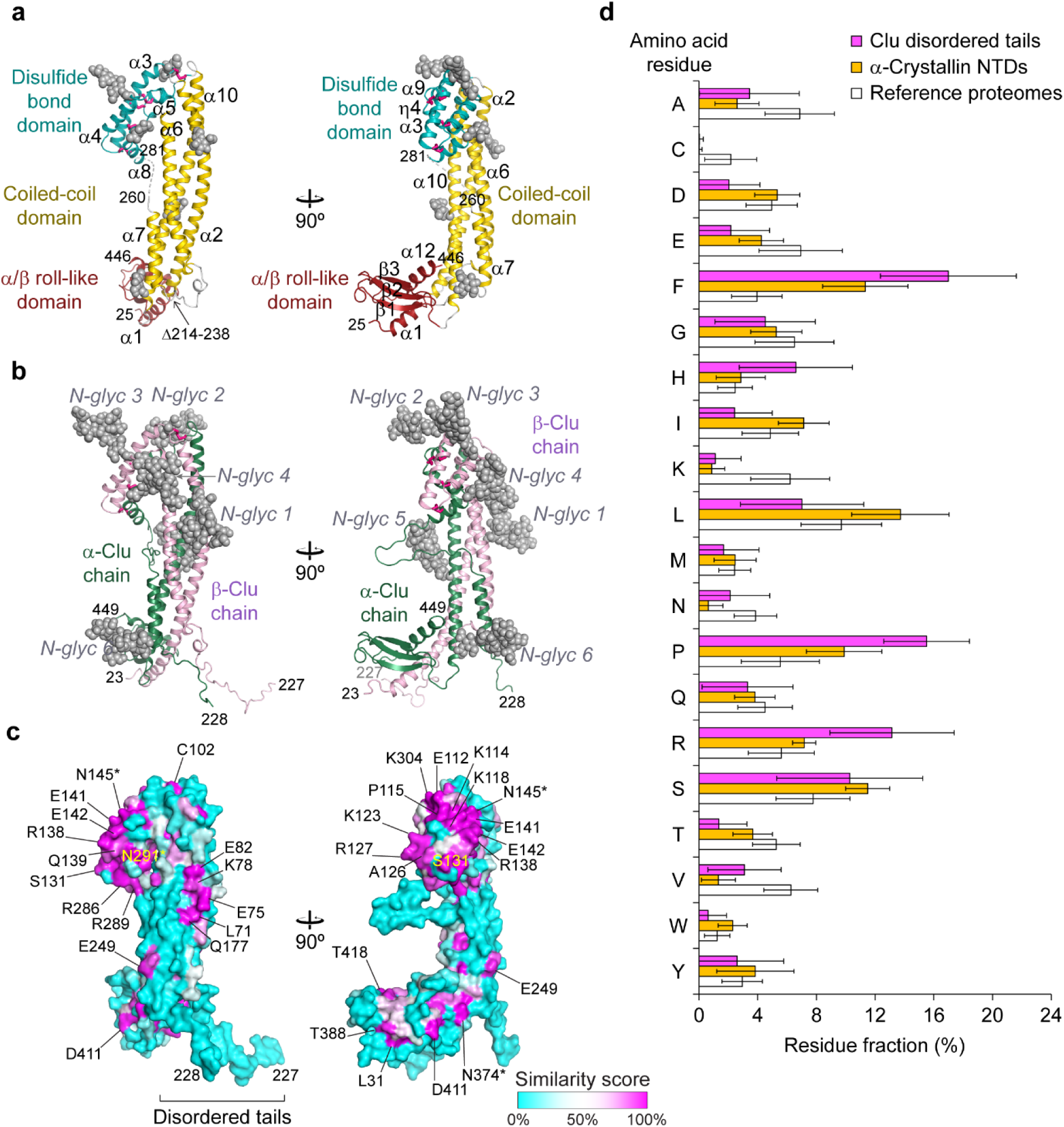
Structural dissection of clusterin chaperone activity. (**a**) Overview of structure-guided mutations in Clu. Clu mutants are named and colored according to their location in disordered tails (TL, blue), disulfide domain (DD, cyan), coiled-coil (CC, orange) and α/β roll-like domain (AB, purple). Amino acid substitutions are listed on the left. The substituted or deleted residues are mapped onto the Clu structural model on the right. Substituted residues are shown in space-filling mode; deleted regions are marked by colored backbone ribbon. (**b**, **c**) Effect of Clu mutations on aggregation of denatured rhodanese (D-Rho). D-Rho was diluted into buffer (final concentration 0.5 µM) containing either no protein, WT-Clu or mutants at molar ratios Clu/D-Rho 1:1 or 3:1. D-Rho aggregation at 25 °C was monitored by turbidity at 320 nm. (b) Representative normalized absorbance traces of D-Rho alone (black), with additional WT-Clu (1:1, dark red; 3:1, light red) and with TL1 (1:1, dark blue; 3:1, light blue) are shown. (c) Bar graph showing the quantification of the absorbance plateau determined by curve fitting in presence of the respective Clu mutant, normalized to D-Rho alone (red and light red dashed lines indicate values for WT-Clu/D-Rho 1:1 and 3:1, respectively). Data represent averages ± SEM (*n* ≥ 3 independent experiments). * p<0.05, ** p<0.01, *** p<0.001 and **** p<0.0001 by one-way ANOVA with Dunnett’s post hoc test comparing Clu mutant/D-Rho 1:1 versus WT-Clu/D-Rho 1:1 and Clu mutant/D-Rho 3:1 to WT-Clu/D-Rho 3:1. (**d**, **e**) Effect of Clu mutations on Aβ(1–42) amyloid formation. Aβ(M1–42) amyloid formation was monitored by thioflavin-T (ThT) fluorescence assay in absence or presence of WT-Clu or mutants (molar ratio Clu/Aβ: 1:300). (d) Representative normalized fluorescence curves in absence (black) or presence of WT-Clu (red) or TL1 (blue). (e) Bar graph showing the relative delay of Aβ aggregation by WT-Clu and mutants determined by the half time of the aggregation plateau (grey and red dashed lines indicate values for Aβ alone and with WT-Clu, respectively). Data represent averages ± SEM (*n* ≥ 3 independent experiments). * p<0.05, *** p<0.001 and **** p<0.0001 by one-way ANOVA with Dunnett’s post hoc test comparing Aβ alone and Clu mutants/Aβ to WT-Clu/Aβ. (**f-h**) Effect of Clu hydrophobic tails on Aβ amyloid formation. (f) Structural model of GFP-TL, a GFP fusion construct carrying the Clu flexible tails at N- and C-termini. (g) Aβ amyloid formation was monitored as in (d) in absence (black) or presence of WT-Clu (red), GFP (grey) or GFP-TL (blue) (molar ratio Clu or GFP/Aβ: 1:300). Representative normalized ThT fluorescence curves are shown. (h) Bar graph showing half times of aggregation compared to Aβ alone (red dashed line). Data represents averages ± SEM (*n* = 3 independent experiments). *** p<0.001 and **** p<0.0001 by one-way ANOVA with Dunnett’s post hoc test comparing Clu or GFP/Aβ to Aβ.

### Structural basis of chaperone function

To identify regions in Clu that mediate the interaction with non-native client proteins, we used rhodanese as a model substrate. Rhodanese (33 kDa) rapidly aggregates upon dilution from denaturant, as detected by turbidity assay (Figure 2b). WT-Clu at a 1:1 and 3:1 molar ratio to denatured rhodanese (D-Rho) efficiently suppressed aggregation at pH 7.2 (Figure 2b) and formed soluble complexes with D-Rho of ∼700 kD, apparently representing D-Rho oligomers bound to multiple molecules of Clu (Supplementary Figure 6a). This ‘holdase’ activity was markedly diminished at pH 5.2, where Clu dimers are absent (Supplementary Figure 1b and 6b).

Strikingly, deletion of residues 214–238 in mutant TL1 (Figure 2a), comprising most of the flexible tail sequences, almost completely abolished the ability of Clu to prevent D-Rho aggregation (Figure 2b, c). To exclude that this loss of chaperone function was due to the single chain nature of TL1, we generated the furin-site mutant TL4, resulting in an otherwise WT single-chain version of Clu (Figure 2a). TL4 showed almost normal holdase activity (Figure 2c), indicating that the loss of aggregation prevention in TL1 is due to the lack of the tail residues.

Similarly, Clu mutants in which aromatic and hydrophobic residues in the β- and α-tails were substituted by Ser (residues 204–227 and 228–238 in TL2 and TL3, respectively, and combined in TL2+3, Figure 2a) showed reduced aggregation prevention to an extent correlating with the number of substitutions (Figure 2c). The only other mutant with a similar aggregation prevention defect for D-Rho was DD1 (Figure 2c), containing multiple substitutions in a conserved surface patch of the DD. Mutants CC3 and CC4 with altered surface clefts at the coiled-coil had intermediate defects (Figure 2c). Thus, the hydrophobic residues in the flexible tails of Clu are critical for chaperone function and cooperate with the DD and adjacent regions in the interaction of Clu with small, soluble aggregates of a non-native protein.

To extend the analysis of Clu chaperone function to a physiologically relevant client, we next monitored the effect of Clu mutants on the nucleation-dependent formation of Aβ(1–42) fibrils. Clu has been shown to be effective at substoichiometric concentrations in delaying Aβ amyloid formation (Beeg et al. 2016, Scheidt, Lapinska et al. 2019). Indeed, WT-Clu at a ratio to Aβ of 1:300 increased the half-time of amyloid formation (Arosio et al. 2015) from ∼1.3 h to ∼4 h (Figure 2d). In contrast, the tail deletion mutant TL1 was almost completely inactive in delaying Aβ aggregation (Figure 2d, e). Mutants TL2, TL3 and TL2+3 retained partial activity (Figure 2e). The deletion mutant CC1, which lacks helix α7 and two short sequence motifs proposed to interfere with Aβ aggregation (Spatharas et al. 2022), was fully active (Figure 2e), and so were mutants CC3, CC4 and DD3 (Figure 2e). However, the single-chain mutant TL4, as well as mutants CC2 and DD2 were partially defective in delaying Aβ amyloid formation, in contrast to their largely unimpaired activity in preventing D-Rho aggregation (Figure 2c, e). Thus, the flexible hydrophobic tails gain full activity in Aβ aggregation prevention only after furin cleavage and function cooperatively with regions outside the tails (Figure 2e). Surprisingly, mutant DD1 was hyperactive in the Aβ aggregation assay (Figure 2e), extending the aggregation half time from ∼4 h for WT-Clu to over 12 h (Supplementary Figure 6c). The DD1 mutations essentially scramble highly conserved residues in a surface patch, thereby apparently enhancing the contribution of this domain to Aβ aggregation prevention, although reducing the interaction with the heterologous substrate D-Rho (Figure 2c). Of note, the DD1 site did not act autonomously, as combining DD1 with the tail deletion TL1 completely abolished Aβ aggregation inhibition (Supplementary Figure 6d). Mutants TL1 and TL2+3 were also defective in slowing amyloid formation of α-synuclein and the repeat domain of tau (TauRD) (Whiten, Cox et al. 2018, Yuste-Checa, Trinkaus et al. 2021) (Supplementary Figure 6e-h).

In order to assess the effect of the flexible tail regions of Clu in isolation, we transplanted the tails onto the chain termini of GFP (GFP-TL) (Figure 2f). GFP itself has no known chaperone function and its N- and C-termini are similar in distance to Clu residues 204 and 238, from which the β- and α-tails emanate. While GFP alone had no effect, GFP-TL was nearly as effective as WT-Clu in delaying amyloid formation (Figure 2g, h). However, GFP-TL was ineffective in suppressing D-Rho aggregation (Supplementary Figure 6i), presumably because the complementary DD region as well as the glycans that confer high Clu solubility are missing.

Taken together, our data establish a critical role of the disordered hydrophobic tails for the interactions of Clu with aggregation-prone client proteins and peptides. While partially effective in aggregation prevention on their own, these regions functionally cooperate with surfaces in the DD. Apparently, mutation of the DD can optimize Clu for prevention of Aβ amyloid formation.

### Role of flexible tails in clusterin uptake

Clu is thought to facilitate cellular uptake of bound substrate proteins via receptor-mediated endocytosis utilizing various surface receptors (Wyatt, Yerbury et al. 2011, Leeb et al. 2014, Yeh, Wang et al. 2016, Itakura, Chiba et al. 2020). We monitored Clu uptake with fluorescently labelled Clu (Clu-A488) in induced pluripotent stem cell-derived neurons (iNeurons) (Supplementary Figure 7a). Uptake of Clu-A488 was competed by unlabeled Clu (Supplementary Figure 7b). Interestingly, the tail mutants TL1, TL2 and TL2+3 exhibited strong uptake defects (∼15–27% residual uptake relative to WT-Clu), with milder defects of TL3 (Figure 3a). Thus, the same structural elements mediating client protein binding are also critical in cellular uptake. Notably, most other mutants also showed mild to intermediate uptake defects including a previously described variant corresponding to DD3 in the present study (Itakura, Chiba et al. 2020), with a ∼55% reduction in uptake (Figure 3a). This may reflect the complexity of Clu binding to a variety of cellular receptors (Wyatt, Yerbury et al. 2011, Leeb, Eresheim and Nimpf 2014, Yeh, Wang et al. 2016, Itakura, Chiba et al. 2020). Thus, multiple regions in Clu, including the flexible tail sequences, are involved in cellular uptake, possibly by mediating interactions with different surface receptors.

**Figure 3:**
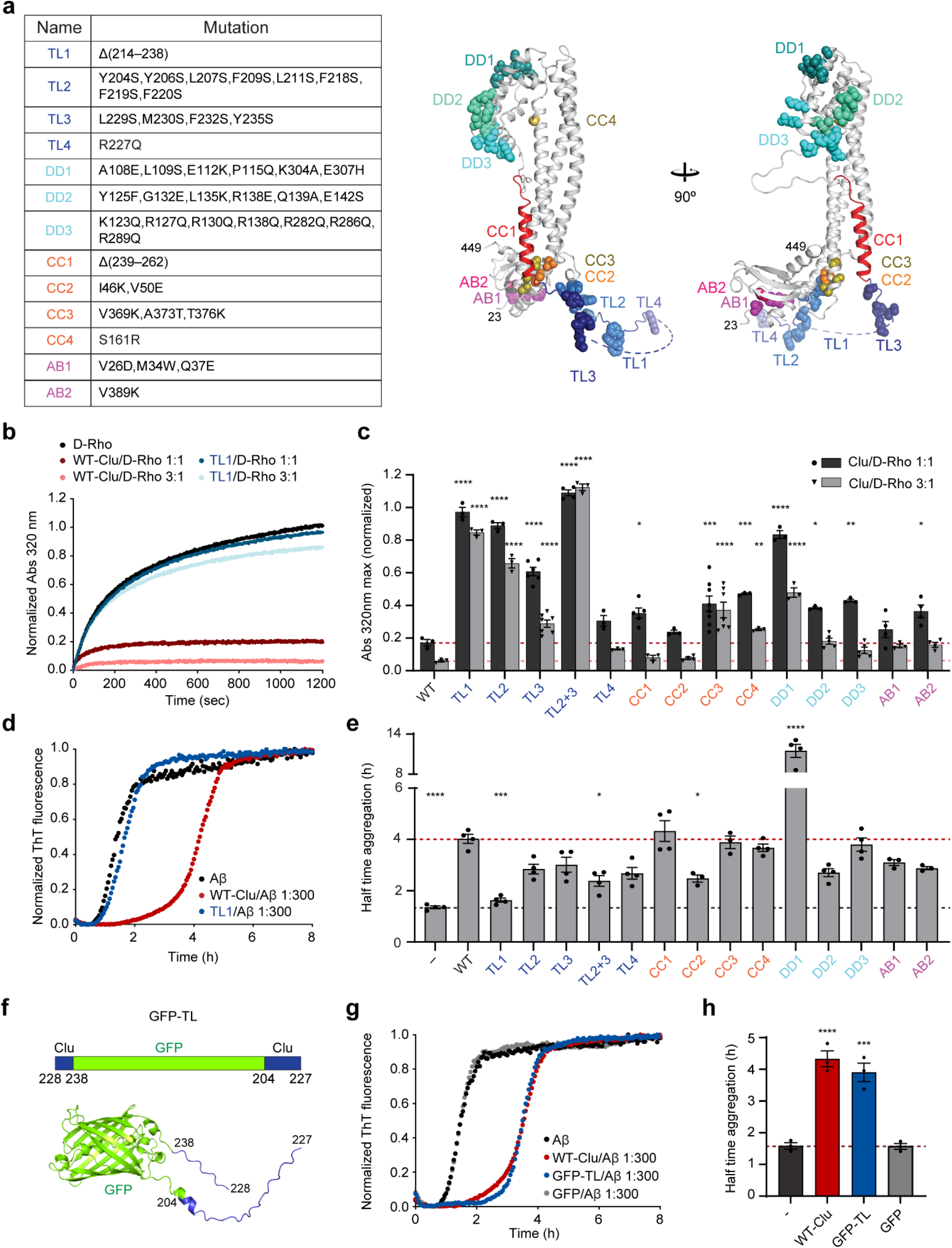
Clusterin cellular uptake and receptor binding. (**a**) Clusterin uptake by neuronal cells. Fluorescently labeled WT-Clu or mutants (Clu-A488) were added to iPSC-derived neurons (iNeurons) and uptake monitored by flow cytometry. Bar graphs represent the percentage of uptake of each mutant relative to WT-Clu (red dashed line at 100% indicates WT-Clu uptake). Data represent averages ± SEM (*n* ≥ 3 independent experiments). * p<0.05, ** p<0.01 and **** p<0.0001 by one-way ANOVA with Dunnett’s post hoc test comparing Clu-A488 mutants to WT-Clu-A488. (**b**) Binding of WT-Clu to VLDLR detected by immune affinity chromatography. WT-Clu and VLDLR ectodomain (VLDLR-ed) with a C-terminal C-tag were incubated for 2 h at equimolar concentration in presence of anti-C-tag affinity resin. WT-Clu alone was used as a background control. After extensive washing, bound proteins were eluted with high-salt buffer and analyzed by SDS-PAGE and immunoblotting. A representative immunoblot is shown (*n* = 3 independent experiments). (**c**) Affinity of WT-Clu and selected mutants to immobilized VLDLR-ed. Microtiter plates were coated with VLDLR-ed and incubated with a series of Clu variant concentrations (50-2000 nM). After extensive washing, bound Clu was quantified by immunodetection. Binding to wells coated with bovine serum albumin (BSA) was used to determine the background signal and subtracted. Individual data points are plotted (*n* =3 independent experiments). Binding curves and dissociation constants (*K*D) for WT-Clu and the mutants TL1 and TL2+3 are shown. n.d., not determined.

We next analyzed the interaction of Clu mutants with very low density lipoprotein receptor (VLDLR), an LDL-type receptor implicated in Clu uptake (Mahon et al. 1999, Leeb, Eresheim and Nimpf 2014) (Supplementary Figure 7c). We expressed and purified the VLDLR ectodomain (VLDLR-ed), which includes the ligand binding domain and a regulatory motif (Rudenko et al. 2002) (Supplementary Figure 7c). Clu interacted directly with VLDLR-ed as shown by VLDLR-ed immunoprecipitation (Figure 3b). We used an enzyme linked immunosorbent assay (ELISA) with immobilized VLDLR-ed to estimate affinities for WT-Clu and mutants. WT-Clu exhibited single-site binding characteristics with a dissociation constant (*K*D) of ∼80 nM (Figure 3c). For comparison, we tested the binding of the receptor-associated protein (RAP) (Supplementary Figure 7c), an ER-resident chaperone known to compete with ligand binding to LDL receptors (Kounnas et al. 1995, Leeb, Eresheim and Nimpf 2014). RAP showed higher affinity (*K*D ∼1.2 nM) (Andersen et al. 2003, Clark et al. 2022) (Supplementary Figure 7d) and competed with Clu for VLDLR-ed binding (Supplementary Figure 7e), suggesting overlapping interaction sites in the VLDLR ligand binding domain (Supplementary Figure 7c).

Binding to VLDLR-ed was disrupted in the Clu tail mutants TL1, TL2 and TL2+3 (Figure 3c and Supplementary Figure 7f), confirming that the tail sequences directly participate in receptor binding. Interestingly, the previously described uptake mutant DD3 (Itakura, Chiba et al. 2020) and the partially overlapping DD2 mutant also exhibited a strongly reduced binding affinity for VLDLR-ed, suggesting the existence of a secondary receptor interaction site in the DD (Supplementary Figure 7g). All other tested Clu mutants, with the exception of the deletion mutant CC1, showed VLDLR-ed affinities close to WT-Clu (Supplementary Figure 7f-h).

In summary the hydrophobic disordered tails of Clu function in receptor-mediated endocytosis. For binding to VLDLR, these sequences cooperate with accessory regions in the DD distinct from the sites contributing to chaperone activity. We suggest that Clu dimers when bound to client protein oligomers or small aggregates retain free tail sequences for receptor association.

### Formation of clusterin-phospholipid particles

Lipoprotein particles containing Clu (also known as ApoJ) are present in human blood plasma and CSF (de Silva, Stuart et al. 1990, Suzuki, Tozuka et al. 2002). To investigate the structural basis of Clu lipid binding, we formed complexes of Clu with the neutral phospholipid 1,2-dimyristoyl-*sn*-glycero-3-phosphocholine (DMPC) (Yeh, Wang et al. 2016, Fernandez-de-Retana et al. 2017). Analysis by native polyacrylamide electrophoresis (native PAGE) revealed the formation of a high molecular weight Clu–lipid complex above a critical molar ratio of Clu/DMPC of ∼1:500 (Figure 4a), similar to the behavior of apolipoproteins ApoA1 and ApoE (Supplementary Figure 8a, b). In contrast, the molecular chaperone Hsc70 did not form complexes with DMPC (Supplementary Figure 8c). The Clu–DMPC complexes fractionated at ∼1–3 MDa in SEC, and negative stain electron microscopy revealed oval disc-shaped particles of ∼19 and ∼26 nm in diameter (Figure 4b and Supplementary Figure 8d). We determined a molar ratio of Clu to DMPC of ∼1:100 in these nanodiscs, corresponding to ∼9 and 16 Clu molecules per nanodisc particle, respectively, suggesting that Clu may associate with both the rim and the surfaces of the nanodiscs. The Clu molecules in nanodiscs are exchangeable, as judged by the ability of excess unlabeled Clu to displace lipid-bound Clu-A488 (Supplementary Figure 8e).

**Figure 4:**
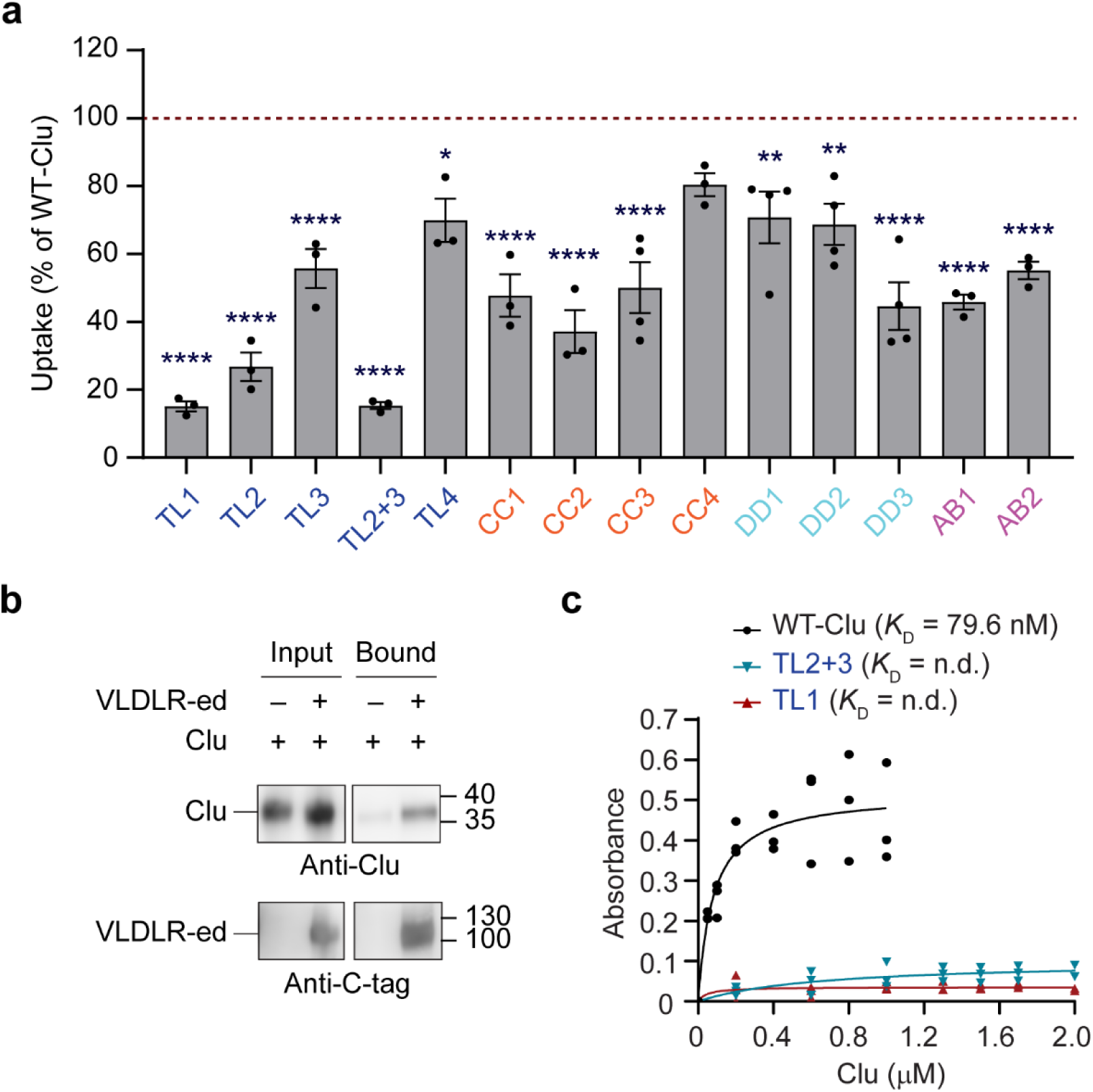
**Probing structure and function of clusterin-lipoprotein complexes**. (**a**) Formation of Clu lipoprotein complexes detected by native PAGE. WT-Clu and DMPC at the indicated molar ratios were cycled three times above (30 °C) and below (18 °C) the melt transition temperature of DMPC (24 °C). WT-Clu alone is shown as a reference. Representative Coomassie blue (protein, left) and Sudan black (lipid, right) stained native gels are shown (*n* = 3 independent experiments). The bands for lipoprotein complex (Clu–DMPC) and free protein (Clu) are indicated. (**b**) Analysis of Clu-lipoprotein complexes by negative stain electron microscopy. A representative micrograph of purified Clu-lipoprotein complexes (Clu–DMPC) is shown (*n* = 3 independent experiments). Side and top views of nanodiscs are visible, indicated by yellow and red arrows, respectively. Scale bar, 100 nm. (**c**) Effect of Clu-lipoprotein complexes on aggregation of denatured rhodanese. Aggregation assays were performed as in Figure 2b with purified Clu-lipoprotein complexes at the indicated molar ratios (shades of blue). Curves for free Clu are shown for comparison (shades of red). Representative light scattering curves are shown (*n* ≥3 independent experiments). Quantification shown in Supplementary Figure 9a. (**d**) Effect of Clu-lipoprotein complexes on Aβ amyloid formation. Aβ amyloid formation was monitored as in Figure 2d in absence (black) or presence of Clu (red), purified Clu-lipoprotein complexes (Clu–DMPC, blue) or DMPC (grey) (molar ratio Clu/Aβ: 1:300 or corresponding amount of DMPC). Representative normalized traces are shown (*n* = 4 independent experiments). Quantification shown in Supplementary Figure 9b. (**e**) Analysis of structural determinants of Clu-lipoprotein complex formation. Formation of lipoprotein complexes with Clu mutant proteins was monitored by native PAGE at a Clu/DMPC ratio of 1:500. Representative Coomassie blue stained native gels are shown (*n* = 3 independent experiments). Quantification shown in Supplementary Figure 9f. (**f**) Lipoprotein complex formation with segments of Clu fused to GFP. GFP fusion proteins with N-terminal α-tail (αTL), Clu helix α7 (H7) and α-tail and helix α7 combined (αTL-H7) were expressed and purified. The domain structure of the constructs is shown on the left. Lipoprotein complex formation was performed as in panel (a) at a 1:1000 molar ratio (construct/DMPC) (*n* = 3 independent experiments). GFP fluorescence of the native PAGE gel was analyzed (middle) and the gel stained with Sudan black (right). The GFP fluorescence signal of the free protein is saturated.

The Clu–DMPC nanodisc preparations were nearly as effective as free Clu in preventing D-Rho aggregation (Figure 4c and Supplementary Figure 9a) and only slightly less active than free Clu in delaying Aβ aggregation (Figure 4d and Supplementary Figure 9b). To determine whether Clu retains chaperone activity in the lipid-bound state, we added nanodiscs to a D-Rho aggregation reaction and then separated them from free Clu by flotation in a density gradient (Supplementary Figure 9c). A fraction of D-Rho was recovered with the Clu–DMPC complexes, indicating that the Clu-containing nanodiscs participate directly in holdase activity (Supplementary Figure 9c). To assess the contribution of the Clu hydrophobic tails to chaperone activity when lipid-bound, we employed limited proteolysis with chymotrypsin, which cleaves after aromatic residues.

Analysis of free Clu by mass spectrometry showed that the β-tail (residues 204–227, mutated region in TL2 mutant) is most sensitive to cleavage, consistent with flexibility and solvent exposure (Supplementary Figure 9d). Notably, incorporation into nanodiscs of WT-Clu or the TL4 single chain mutant had little effect on cleavage by chymotrypsin, suggesting solvent exposure of this region in the nanodiscs as well (Supplementary Figure 9d, e). Thus, the nanodisc particles are chaperone active, likely employing the flexible tails of Clu for client binding.

Surprisingly, lipoprotein formation was reduced by ∼80-90% in mutants deleting or changing the tail sequences (TL1, TL2, TL3 and TL2+3) (Figure 4e and Supplementary Figure 9f). It was essentially eliminated with mutant TL2+3, in which all hydrophobic residues in the tails are substituted by Ser (Figure 4e and Supplementary Figure 9f), while single-chain Clu (TL4) showed normal lipid binding (Figure 4e and Supplementary Figure 9f). Interestingly, lipid binding was also abolished upon deletion of the amphipathic helix α7 (mutant CC1) that follows on the α-tail (mutant TL3) (Figure 4e and Supplementary Figure 9f). Of note, in lipoprotein particles containing ApoE or ApoA1, pairs of (albeit longer) amphipathic helices shield the hydrophobic rim of phospholipid discs (Bibow et al. 2017, Henry et al. 2018). Using GFP fusion constructs, we found that isolated α-tail and helix α7 in combination (residues 228–262; αTL-H7) conferred detectable lipid binding activity (Figure 4f), consistent with the mutational data (Figure 4e and Supplementary Figure 9f), while constructs containing helix α7 (H7) or the α-tail alone (αTL) were not sufficient (Figure 4f). αTL-H7 and DMPC formed disc-shaped particles similar to full-length Clu, with small densities – likely GFP moieties – lining the rims (Supplementary Figure 9g). In the context of the lipoprotein complex, the α-tail might extend helix α7. The Clu mutants DD2 and DD3 also showed markedly diminished incorporation into lipid particles (Figure 4e and Supplementary Figure 9f). Nanodisc formation thus depends on multiple structural elements of Clu, including the flexible tails and the amphipathic helix α7, with the DD having an accessory role.

These data are consistent with a model in which the hydrophobic tail sequences initiate lipid binding, followed by undocking of helix α7 from the coiled-coil and association with the nanodisc rim. The hydrophobic tails, while partitioning into the phospholipid bilayer, remain dynamic and accessible for chaperoning misfolded client proteins.

## Conclusion

Our structure-based analysis of clusterin function provides insight into the mechanism of action of this medically important extracellular chaperone and apolipoprotein (Foster et al. 2019, Wilson, Satapathy and Vendruscolo 2023, Mamun et al. 2024). The crystal structure of human Clu revealed a discontinuous three-domain architecture consisting of a coiled-coil bundle flanked by a conserved helical domain stabilized by disulfide bonds and an α/β roll-like domain (Figure 1a). Surprisingly, we found two disordered hydrophobic tails, generated by cleavage of the central region of the Clu precursor, to be critical for multiple functionalities, including chaperone activity, binding to cell surface receptors, and lipoprotein complex formation (Figure 5a).

**Figure 5:**
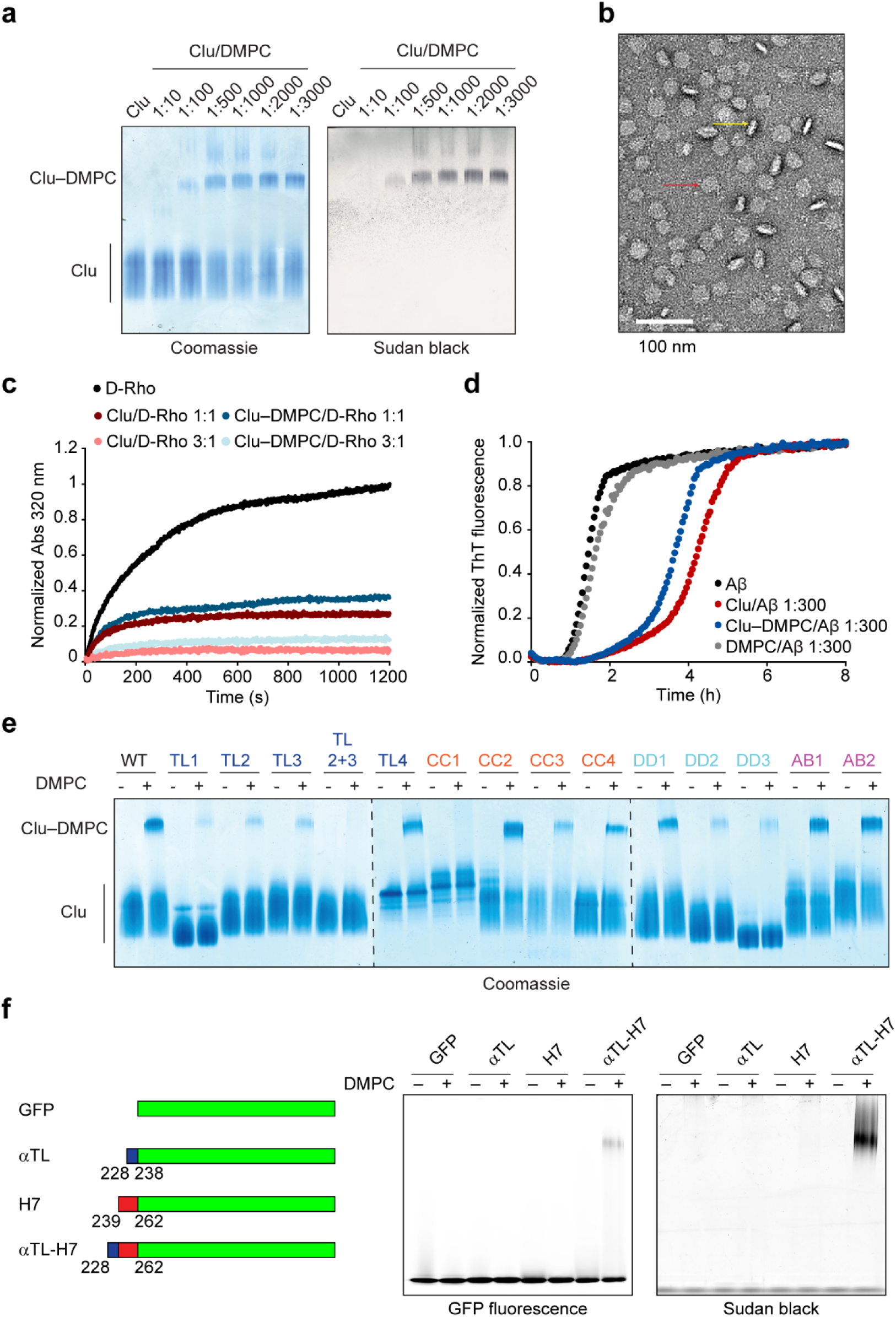
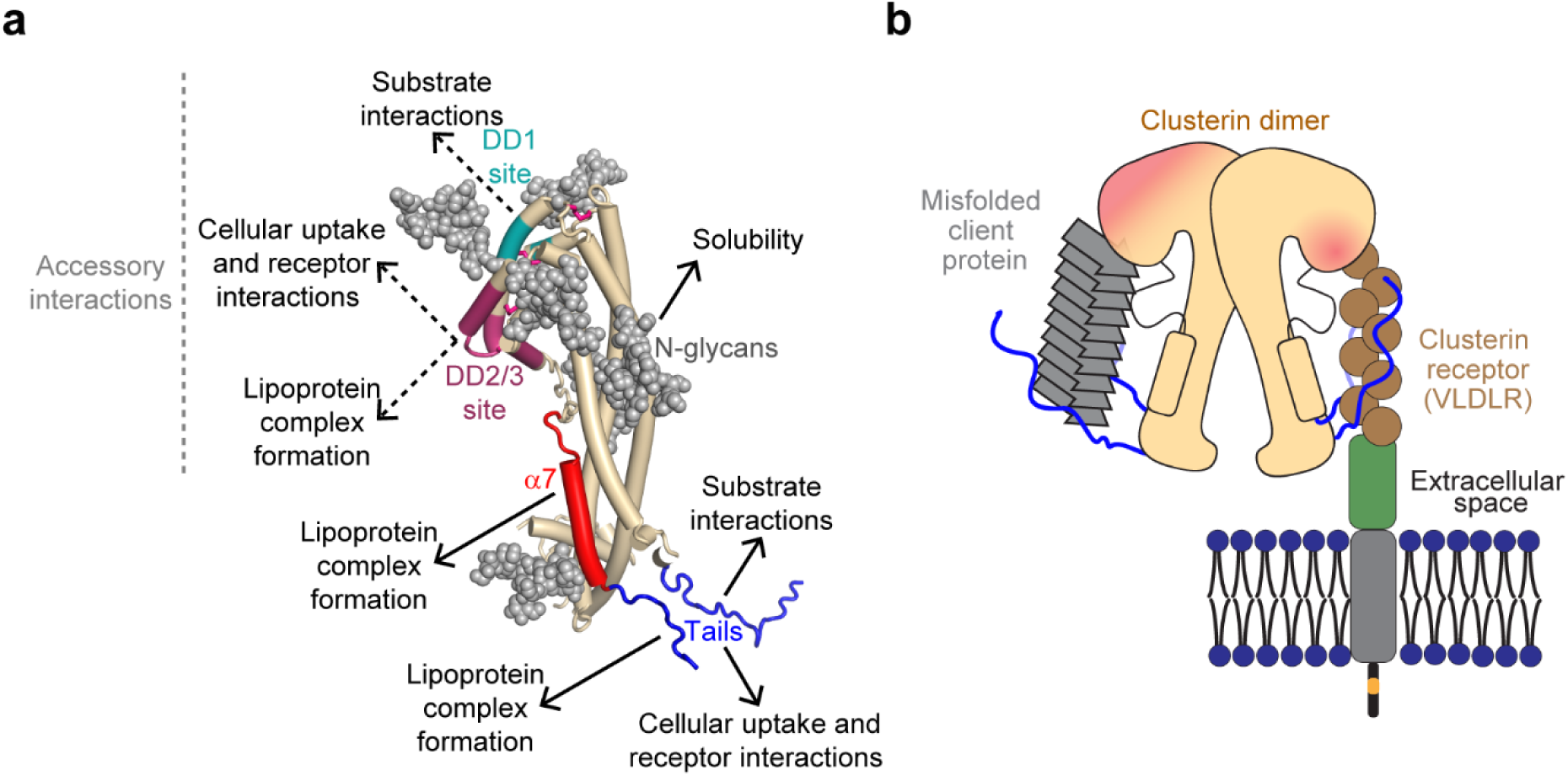
Functional assignment of clusterin structural elements. (**a**) Structural determinants for the functions of extracellular Clu. The respective main and accessory regions for substrate interactions, cellular uptake and lipoprotein complex formation were mapped onto a simplified structure model of Clu. Helices are represented as tubes. Disordered tails (blue) and helix α7 (red) as well as the DD1 (teal) and DD2/3 (purple) sites are indicated. N-glycans (gray) are shown in space-filling mode. (**b**) Model for cellular uptake of Clu bound to substrate. The disordered tails of Clu presumably interact with the ligand binding repeats of LDLR type receptors (brown circles, Supplementary Figure 7c). Formation of Clu dimers or binding of multiple Clu molecules to small aggregates (gray) might enable simultaneous interaction with client protein and cell surface receptors.

The hydrophobic tails of Clu resemble the N-terminal substrate binding regions of intracellular sHsp chaperones in terms of amino acid composition and biophysical properties (Woods, Ulmer et al. 2023), suggesting convergent evolution of chaperone activity in the phylogenetically younger Clu. The enrichment of these sequences in Clu with Phe, His and Arg residues might enable π-stacking interactions with non-native substrate proteins. Indeed, deletion of the tails or replacement of their hydrophobic residues with serine resulted in a loss of chaperone function in stabilizing non-native proteins in solution and inhibiting amyloid formation by Aβ(1–42), α-synuclein and tau. Considering the extensive modification of Clu with hydrophilic N-glycans, Clu would be highly efficient in solubilizing aggregation-prone client proteins, rendering them available for cellular uptake and degradation. Clu dimer formation at physiological pH and the interaction of multiple Clu molecules with small protein aggregates presumably ensure that hydrophobic tail sequences are available for both chaperone function and receptor binding (Figure 5b). Notably, the hydrophobic tails of lipid-bound Clu remain dynamic and accessible, explaining why Clu lipoprotein complexes are chaperone active (Calero et al. 1999).

The tail sequences of Clu cooperate in aggregation prevention with a highly conserved region in the Clu disulfide domain (DD). We found this effect to be relevant in the ability of Clu to delay the nucleation dependent aggregation of Aβ, where Clu is effective at substoichiometric concentrations (Beeg, Stravalaci et al. 2016, Scheidt, Lapinska et al. 2019). Interestingly, the DD1 variant of Clu, containing multiple mutations in a highly conserved DD surface patch, was hyperactive in Aβ aggregation prevention, in a manner dependent on the hydrophobic tails. It would appear, therefore, that the chaperone activity of Clu has not been optimized in evolution to suppress the formation of Aβ amyloid, allowing the DD to function more broadly in interacting with non-native client proteins as well as with membrane receptors. Indeed, the DD1 mutant was markedly impaired in D-Rho aggregation prevention, while the DD2 and DD3 mutants showed reduced VLDLR binding. Our findings thus demonstrate that Clu can be adapted to specific client proteins. It will be of interest to explore the therapeutic potential of the DD1 mutant and other Clu variants in mouse models of AD.

## Methods

### Plasmids

The pB-T-PAF-Clu plasmid used for wildtype clusterin (WT-Clu) expression and purification was constructed by amplifying the coding region of pB-T-PAF-CluStrep (Yuste-Checa, Trinkaus et al. 2021) via PCR. The PCR product was then digested with NheI and NotI and subcloned into the pB-T-PAF vector. All pB-T-PAF plasmids encoding Clu variants are based on this vector. pB-T-PAF-Clu-TL1 was obtained by PCR-amplification of the entire plasmid with the mutagenic primers using Herculase-II (Agilent), followed by ligation with the KLD enzyme mix (NEB). pB-T-PAF-Clu-CC1 and pB-T-PAF-Clu-TL4 were obtained by separate PCR-amplification of 5’- and 3’-Clu fragments with the mutagenic primers, followed by digestion with NheI and NotI and simultaneous subcloning into pB-T-PAF. pB-T-PAF-Clu-TL2, TL3, TL2+3, CC2, CC3, DD1, DD2, DD3, AB1 and AB2, were obtained by plasmid digestion with NheI/NotI or PCR-amplification of the plasmid with primers encoding overlapping regions with the corresponding synthesized gene fragment including the mutations (Twist Bioscience) and assembly with the NEBuilder HiFi DNA Assembly Master Mix (New England Biolabs). pB-T-PAF-Clu-CC4 was obtained by mutagenesis using the Q5 Site-Directed Mutagenesis Kit (New England Biolabs). pB-T-PAF-Clu-DD1+TL1 was obtained by PCR amplification of pB-T-PAF-Clu-DD1 plasmid to delete the TL1 region and assembly with the NEBuilder HiFi DNA Assembly Master Mix (New England Biolabs).

pB-T-PAF-VLDLR(1-797)Δ(751-779)-C-tag was obtained by PCR amplification of VLDLR from human placenta cDNA (Edge Biosystems) using Herculase-II Fusion DNA polymerase (Agilent Technologies), followed by Gibson assembly with pB-T-PAF using NEBuilder HiFi DNA Assembly Master Mix (New England Biolabs).

pET-Sac-Abeta(M1-42) plasmid for amyloid-β (M1-42) expression and purification was a gift from Dominic Walsh (Addgene plasmid # 71875) (Walsh et al. 2009). pQTEV-LRPAP1 plasmid for RAP expression and purification was a gift from Konrad Büssow (Addgene plasmid # 31327) (Büssow et al. 2005). The signal peptide was removed (amino acids 1-35) by mutagenesis using the Q5 Site-Directed Mutagenesis Kit (New England Biolabs).

pHUE-eGFP-Clu-tail was generated by restriction and insertion cloning with a DNA fragment encoding eGFP-Clu-tail amplified from pEGFP-C2 (Clontech) by nested PCR using primers encoding Clu(228-238) and Clu(204-227) and introducing SacII and HindIII restriction sites and the SacII/HindIII restriction fragment of plasmid pHUE (Catanzariti et al. 2004).

pHUE-Clu-αTL-H7-eGFP, pHUE-Clu-H7-eGFP and pHUE-Clu-αTL-eGFP vectors were obtained by PCR amplification and assembly with the pHUE-eGFP-Clu-tail vector backbone digested with SacII/HindIII using the NEBuilder HiFi DNA Assembly Master Mix (New England Biolabs).

All newly made plasmids were deposited in Addgene (Supplementary Table 2). <u>Cell lines</u> HEK293-EBNA (HEK293E) suspension cell lines (Durocher et al. 2002) stably expressing the recombinant Clu and VLDLR protein constructs (Supplementary Table 2) were generated using a piggyBac transposon-based expression system (Li et al. 2013) employing the respective pB-T-PAF-Clu plasmids. Newly generated cell lines were registered at Cellosaurus.org (RRID available in Supplementary Table 2).

Induced pluripotent stem cell (iPSC) line HPSI0214i-kucg_2 (RRID:CVCL_AE60) was purchased from UK Health Security Agency (#77650065, supplied by HipSci). iPSCs were maintained at 37 °C and 5% CO2 in mTeSR or mTeSR plus medium (Stem Cell Technologies) on Geltrex-coated (Thermo Fisher Scientific) cell culture plates. Cells were split when confluent using ReLeSR (Stem Cell Technologies). Quality control tests are specified in Supplementary Table 3.

Neural progenitor cells (NPCs) were generated using the STEMdif SMADi Neural Induction Kit (Stem Cell Technologies) following the monolayer protocol. NPCs were frozen in STEMdif Neural Induction Medium + SMADi with 10% DMSO.

iPSC derived-forebrain-type neurons were generated from the NPCs described above. NPCs were thawed in STEMdif Neural Induction Medium + SMADi on 0.01% Poly-L-ornithine/10-20 µg ml^−1^ laminin-coated (MERCK) cell culture plates and differentiated using the STEMdiff Forebrain Neuron Differentiation Kit (Stem Cell Technologies) followed by the STEMdiff Forebrain Neuron Maturation Kit (Stem Cells Technologies). STEMdiff Forebrain Neuron Maturation medium was supplemented for 6 days with 5 µM 5-fluorouracil/uridine (Merck) to stop growth of non-differentiated cells. After 8 days in STEMdiff Forebrain Neuron Maturation medium (half medium change every other day), iPSC-derived neurons were maintained in Neurobasal Plus medium (Thermo Fisher Scientific) supplemented with B-27 Plus Supplement 1x (Thermo Fisher Scientific), 0.5 mM GlutaMAX (Thermo Fisher Scientific) and 100 U ml^−1^ penicillin, 100 µg ml^−1^ streptomycin sulfate (Thermo Fisher Scientific).

### Gel electrophoresis and immunoblotting

For sodium dodecyl sulfate-polyacrylamide gel electrophoresis (SDS-PAGE), protein samples were boiled in SDS-PAGE sample buffer for 5 min and separated by electrophoresis on NuPAGE 4–12% Bis-Tris SDS gels (Thermo Fisher Scientific) using NuPAGE MES SDS running buffer (Thermo Fisher Scientific) at 140 V. For native PAGE, the samples were mixed with NativePAGE Sample Buffer (4X) (Thermo Fisher Scientific) and separated by electrophoresis on NativePAGE 3–12% Bis-Tris SDS gels (Thermo Fisher Scientific) using NativePAGE running buffer (Thermo Fisher Scientific) at 140 V. Proteins were transferred at 70 V for 2 h onto a nitrocellulose membrane (GE Healthcare) using a wet electroblotting system (Bio-Rad). Alternatively, the Power Blotter XL (Invitrogen) with Select transfer stacks nitrocellulose (Invitrogen) was used. Membranes were blocked for at least 1 h with Tris-buffered saline (TBS) containing 0.05% Tween 20 (0.05% TBS-Tween) and 5% low fat milk or 3% bovine serum albumin. Immunodetection was performed using mouse monoclonal Clu-α antibody (Santa Cruz Biotechnology, sc-5289, 1/1000 dilution), rabbit anti-rhodanese (in-house, 1/5000 dilution) and CaptureSelect biotin anti-C-tag conjugate (Thermo Fisher Scientific, 7103252100, 1/2000 dilution). Conjugated goat-anti mouse immunoglobulin G (IgG)-horseradish peroxidase (HRP) (Merck, A4416), goat-anti rabbit IgG-HRP (Merck, A9169) and Streptavidin-HRP (Pierce, 21130, 1/10000 dilution) were used as secondary antibodies.

Immobilon Forte Western HRP substrate (Merck) was used for detection. Full scan blots are provided in the Source Data.

### Protein expression and purification

All protein purification steps were performed at 4 °C unless otherwise indicated. Protein concentrations in the final preparations were determined by absorbance at 280 nm using absorbance coefficients calculated from the protein sequence with the program ProtParam unless otherwise noted. Purified protein samples were concentrated by ultrafiltration and snap-frozen in liquid nitrogen for storage at –70°C.

#### Clusterin constructs

Clu constructs were expressed and secreted by the respective HEK293E stable cell lines cultured in FreeStyle 293 Expression Medium (Thermo Fisher Scientific) for 4 days. For expression of Clu constructs containing oligo-mannose N-glycans, the α-mannosidase-I inhibitor kifunensine (MedChemExpress) dissolved in water was added to the medium (16 μM final concentration).

The conditioned medium was then separated from the cells by centrifugation. For chromatographic purification, 200 ml medium were first dialyzed against wash buffer (20 mM Na acetate pH 5.0). After removal of precipitate by centrifugation, the supernatant was passed over a HiTrap SP XL cation exchange column (Cytiva, 5x 5 ml volume). The column was washed with 5 column volumes (CV) wash buffer. For protein elution, a 0-500 mM NaCl gradient in wash buffer was applied. Clu-containing fractions were further purified by SEC on a HiLoad 26/600 Superdex-200 (Cytiva) in 20 mM Na acetate pH 5.0, 100 mM NaCl, 1 mM EDTA. Fractions containing pure, monomeric Clu were merged. A detailed protocol is available here: dx.doi.org/10.17504/protocols.io.bvvkn64w. Here we omitted the denaturing wash since hardly any contaminants were observed.

#### VLDLR-ed

VLDLR(28-797)Δ(751-779)-C-tag (VLDLR-ed, derived from Uniprot isoform sequence P98155-2) was expressed and secreted by HEK293-VLDLR(1-797)Δ(751-779)-C-tag cells cultured in FreeStyle 293 expression medium (Thermo Fisher Scientific) for 4 days. The conditioned medium was then separated from the cells by centrifugation. For chromatographic purification, the medium was first dialyzed against binding buffer (20 mM Tris-HCl pH 7.2, 100 mM NaCl, 0.5 mM CaCl2) overnight. VLDLR-ed in the dialysate was purified by affinity chromatography using CaptureSelect C-tag affinity matrix (Thermo Scientific) using an elution buffer containing 20 mM Tris HCl pH 7,0, 2 M MgCl2 and 2 mM CaCl2. Finally, the protein buffer was exchanged with a NAP-25 desalting column (Cytiva) equilibrated in binding buffer. A detailed protocol is available here: dx.doi.org/10.17504/protocols.io.j8nlkoqw1v5r/v1.

#### eGFP-Clu constructs

eGFP-Clu constructs were expressed as a C-terminal His6-ubiquitin fusion protein in *E. coli* Bl21-CodonPlus (DE3)-RIL cells (Agilent) transformed with the corresponding plasmid and cultured in 4 l LB medium overnight at 18 °C with 0.25 mM IPTG. After lysis of the cells by ultrasonication on ice in PBS buffer containing 20 mM imidazole, 1 mM phenylmethylsulfonyl fluoride (PMSF) and Complete EDTA-free protease inhibitor cocktail (Roche), the clarified supernatant was subjected to affinity chromatography on Ni-chelating Sepharose (Cytiva 17-0575-01). Green fluorescent elution fractions were merged and digested with His-tagged Usp2 during 4 h on ice. After buffer exchange into PBS buffer containing 20 mM imidazole using a HiPrep 26/10 desalting column (Cytiva 17-5087-01), a second affinity chromatography on Ni-chelating Sepharose (Cytiva 17-0575-01) was performed. The green-fluorescent fractions of unbound protein were pooled and concentrated by ultrafiltration, and the concentrate subjected to SEC on a HiLoad 16/600 Superdex-200 (Cytiva 28-9893-35) column equilibrated with PBS. Concentrations were determined by absorbance at 488 nm using an absorbance coefficient of 61,000 M^−1^ cm^−1^. A detailed protocol is available here: dx.doi.org/10.17504/protocols.io.n2bvj3w55lk5/v1.

#### TauRD

TauRD (Tau residues 244-371, C291A/P301L/C322A/V337M) was expressed and purified as described (Yuste-Checa, Trinkaus et al. 2021). A detailed protocol is available here: dx.doi.org/10.17504/protocols.io.x54v9p6p1g3e/v1.

#### α-Synuclein(A53T)

α-Synuclein(A53T) was expressed and purified as described (Yuste-Checa, Trinkaus et al. 2021). A detailed protocol is available here: dx.doi.org/10.17504/protocols.io.btynnpve.

#### Hsc70

Hsc70 was expressed and purified as described (Schneider et al. 2021).

#### RAP

RAP was expressed as N-terminal His7-TEV site fusion protein in *E. coli* BL21(DE3) cells transformed with the pQTEV-LRPAP1 (without signal peptide) plasmids via IPTG induction. The cell pellet from 6 l culture was resuspended in 200 ml lysis buffer (50 mM Tris-HCl pH 8.0, 300 mM NaCl, 10 mM imidazole) supplemented with 1 mg ml^−1^ lysozyme, Complete EDTA-free protease inhibitor cocktail (Merck) and Sm DNase 50 U ml^−1^, and incubated while gently shaking at 4 °C for 30 min. Cells were lysed by ultrasonication, and the lysate was cleared by centrifugation (1 h, 40,000 × g at 4 °C). The supernatant was loaded onto a Ni-NTA column equilibrated with lysis buffer. His7-TEV-RAP was eluted with elution buffer (50 mM Tris-HCl pH 8.0, 300 mM NaCl, 250 mM imidazole). The eluted fractions were collected and the salt concentration was reduced using a HiPrep 26/10 desalting column with 50 mM Tris-HCl pH 8.0, 10 mM NaCl buffer. The eluted protein was then incubated with 20 ml TEV mg^−1^ protein, 10% glycerol, 1 mM DTT, 0.25 mM EDTA overnight at 4 °C in order to cleave the His7-TEV tag.

The cleavage mixture was applied to a Ni-NTA column equilibrated with 50 mM Tris-HCl pH 8.0, 10 mM NaCl buffer in order to remove the His7-TEV tag and the TEV protease. The flow-through containing RAP was collected and loaded onto a Superdex 200 column equilibrated with PBS pH 7.4. Fractions containing pure protein were combined, concentrated by ultrafiltration using Vivaspin MWCO 10,000 (GE Healthcare), aliquoted and flash-frozen in liquid nitrogen for storage at −70 °C. A detailed protocol is available here: dx.doi.org/10.17504/protocols.io.rm7vzxpb2gx1/v1.

#### Aβ

Aβ (M1-42) was purified as described (Linse 2020) with modifications. Aβ (M1-42) was expressed in *E. coli* BL21(DE3) cells transformed with the pET-Sac-Abeta(M1-42) plasmid via IPTG induction. The cell pellet from a 3 l culture was resuspended in 80 ml TE pH 7.5 buffer (10 mM Tris-HCl pH 7.5, 1 mM EDTA) supplemented with Complete EDTA-free protease inhibitor cocktail (Merck) and 2.5 U ml^−1^ benzonase until sample was homogenous. Cells were lysed by sonication, and the lysate cleared by centrifugation (10 min, 18,000 x g at 4 °C). The pellet was resuspended in 50 ml TE pH 7.5 until the sample was homogenous, sonicated and centrifuged (10 min, 18,000 x g at 4 °C). This step was repeated twice. The pellet was then resuspended in 40 ml TE pH 9.5 until sample was homogenous, sonicated and centrifuged (10 min, 18,000 x g at 4 °C). The supernatant was collected, filtered through a 0.45 μm pore size filter and pH was adjusted to 8.5 using 1 M HCl. The sample was then loaded onto a diethylaminoethyl (DEAE) cellulose column equilibrated with TE pH 8.5. The column was washed with TE pH 8.5 10 mM NaCl and eluted with TE pH 8.5 50 mM NaCl. Fractions containing pure Aβ peptide were combined and buffer was exchanged to TE pH 8.5 using a desalting column. 100 ml elution from the desalting column were frozen in liquid N2 while rotating and lyophilized. The lyophilized sample was resuspended in 15 ml GE buffer pH 8.5 (6 M guanidinium hydrochloride, 20 mM sodium phosphate) and loaded onto a Superdex 75 SEC column equilibrated with AE buffer pH 8.5 (20 mM ammonium acetate, 0.2 mM EDTA). Fractions containing monomeric Aβ were combined and peptide concentration was determined by absorbance at 205 nm. 500 µg aliquots were flash-frozen, lyophilized and stored at −70 °C.

### Removal of N-glycans with PNGase F under denaturing conditions

This procedure is essentially identical to New England Biolabs protocol for PNGase F (https://www.neb.com/en/protocols/2014/07/31/pngase-f-protocol). The Clu variant of interest at 10 μM in Glycoprotein Denaturing Buffer (0.5% SDS, 40 mM DTT) (total volume, 20 μl) was heated to 95 °C for 10 min, followed by cooling on ice. Subsequently, the sample was split 1:1 and to each half 10 μl 1% NP40 and 2X GlycoBuffer (100 mM Na-phosphate pH 7.5) either with or without 0.17 U GST-PNGase F (MPIB Core Facility) were added, followed by incubation at 37 °C for 1 h. The samples were analyzed by SDS-PAGE and Coomassie staining.

### Hydrogen/deuterium exchange–mass spectrometry

#### Sample preparation

Clu was prepared at 200 µM in HDX buffer (20 mM Na-acetate pH 5.0, 100 mM NaCl, 1 mM EDTA, 1 mM tris(2-carboxyethyl)phosphine (TCEP)). To initiate the deuterium exchange reaction, 2.2 µl protein was added to 27.8 µl deuteration buffer (HDX buffer prepared in D2O) and incubated for different times (10, 100 or 1000 s) at 25 °C before quenching the reaction by addition of 79 µl ice cold quench buffer (100 mM sodium phosphate pH 2.4, 20 mM TCEP, 7 M guanidine-HCl). Reactions were incubated 15 min before addition of 109.1 µl sodium phosphate pH 2.4, resulting in a final pH of 2.5 to 2.6.

#### Peptide mass analysis and data processing

Quenched samples were injected into a Waters ACQUITY UPLC M-class instrument with H/DX via a 50 µl sample loop. Proteins were digested using an Enzymate BEH-pepsin column (Waters) at a flow rate of 100 µl min^−1^ and temperature of 20 °C. Peptides were trapped and desalted for 3 min at 100 µl min^−1^ before transfer to a 1.0 x 100 mm ACQUITY UPLC peptide CSH C18 column (Waters) held at 0 °C. Peptides were eluted over 20 min with an 8-40% acetonitrile gradient in 0.1% formic acid, pH 2.5. Between injections, the analytical column was washed using two repeating sawtooth gradients and equilibrated at 8% acetonitrile. Mass analysis was performed on a Waters Synapt G2Si. T-wave ion mobility was used as an orthogonal peptide separation step between the UPLC and mass spectrometer (Iacob et al. 2008). Ion guide settings were adjusted to minimize gas-phase back exchange as described (Guttman et al. 2016). Peptides were identified by analyzing MSE data for 4-5 undeuterated control experiments using PLGS (Waters). Mass spectra were processed in DynamX (Waters, https://www.waters.com/) and peak selection was manually verified for all peptides. All experiments were performed under identical conditions. Deuterium levels were therefore not corrected for back exchange and are reported as relative (Wales and Engen 2006). Experiments were performed in triplicate.

### Circular dichroism spectroscopy

Far-UV CD spectra as well as thermal transitions of proteins were measured with a Jasco J-715 spectrometer equipped with a Peltier-thermostat using 0.1 cm cuvettes. Wavelength scans were recorded at 20 °C, temperature scans at the indicated wavelength applying a temperature gradient of 60 °C h^−1^. The proteins were analyzed at the indicated concentrations in a buffer containing 50 mM potassium phosphate pH 7.0. To estimate the WT-Clu properties at pH 5, a buffer containing 50 mM potassium phosphate pH 5.0 was used.

### Analytical SEC

Purified Clu constructs at a concentration of 10 μM were equilibrated on ice overnight or longer with buffers containing 100 mM NaCl and 1 mM EDTA and either 20 mM Na-acetate pH 5.0, 20 mM MES-NaOH pH 6.5, 20 mM HEPES-NaOH pH 7.5 or Tris-HCl pH 8.5 in a total volume of 60 μl and then analyzed by size exclusion chromatography (SEC) on a Superdex-200 Increase 3.2/300 column (Cytiva) using an NGC chromatography system (Bio-Rad). The system was at room temperature (RT) and 0.05 ml min^−1^ flow rate was used. The injection volume was 50 μl. The runs were recorded at 280 nm wavelength.

For separation of soluble aggregates of Clu with denatured rhodanese and Clu–DMPC lipoprotein complexes by SEC, an Ettan chromatography system (GE Healthcare) equipped with a Superose 6 Increase 3.2/300 column (Cytiva) and PBS running buffer at RT and 0.05 ml min^−1^ flow rate was used.

### Crystallization

Crystals of Clu mutant Clu-Δ(214–238) (TL1) with oligo-mannose N-glycans were obtained with the help of the Max Planck Institute of Biochemistry (MPIB) Crystallization Facility by the sitting-drop vapor diffusion method using the Classics (Qiagen, Hilden, Germany), Index

(Hampton Research, Aliso Viejo, CA) and JCSG-plus (Molecular Dimensions, Sheffield, UK) crystallization screens at 4 °C by mixing 200 nl sample with 200 nl reservoir.

Crystal form I of space group *P*21 was obtained with 30% PEG-4000, 0.2 M ammonium acetate and 0.1 M Na-citrate pH 5.6 (Classics condition H3), 0.05 M ammonium sulfate, 0.05 M BIS-TRIS pH 6.5, 30% v/v Pentaerythritol ethoxylate (15/4 EO/OH) (Index condition E9) and 0.2 M lithium sulfate monohydrate, 0.1 M BIS-TRIS pH 6.5, 25% PEG-3350 (Index condition G3) as precipitant. For cryoprotection, precipitants additionally containing 7.5% and 15% glycerol were prepared. The crystals were serially incubated with these buffers for each 20 min, respectively, before flash-cooling in liquid nitrogen. Diffraction data from the first condition were used for model building and refinement.

Crystal form II of space group *C*2 was obtained with 24% PEG-1500 and 15% glycerol (JCSG-plus condition D1). These crystals were directly flash-cooled in liquid nitrogen.

### Structure solution and refinement

X-ray diffraction data were collected at 100 K and a wavelength of 0.7749 Å by the oscillation method at beamline ID23-1 at the European Synchrotron Radiation Facility (ESRF), Grenoble, France.

The data were integrated and scaled with XDS (Version Feb 5, 2021, https://xds.mr.mpg.de/) (Kabsch 2010). The programs Pointless 1.12.4 (Evans 2006), Aimless 0.7.4 (Evans and Murshudov 2013) and Ctruncate 1.17.29 (French and Wilson 1978), as implemented in the CCP4i graphical user interface (Version 7.1.010, https://www.ccp4.ac.uk/) (Potterton et al. 2003), were used for data reduction. Partial datasets collected from different portions of the crystals were merged. The X-ray diffraction was strongly anisotropic, and an overall I/σ(I) of 1.5 in the outer resolution shell was used as resolution cut-off criterion.

The crystal structure of Clu mutant TL1 (Clu-Δ(214–238)) in space group *P*21 was solved by molecular replacement with Molrep 11.4.06 (CCP4i interface 7.1.010) (Vagin and Isupov 2001) using a truncated version of the Alphafold2 model for human clusterin (https://alphafold.ebi.ac.uk/entry/P10909) (Jumper et al. 2021, Varadi et al. 2022) in which low-confidence regions were trimmed away. WinCoot 0.9.4.1 (http://bernhardcl.github.io/coot) was employed for manual model building (Emsley and Cowtan 2004). The model was refined with Refmac5 5.8.0267 (CCP4i interface 7.1.010) (Murshudov et al. 2011). The final refinement was performed with Phenix.refine 1.19.2-4158 (http://www.phenix-online.org/) (Liebschner et al. 2019). Residues facing solvent channels with disordered side-chains were truncated after C-β. The final model contains one copy of Clu mutant TL1 per asymmetric unit, with residues 23–24, 261–280 and 447–449 missing for lack of discernible electron density, probably because of disorder. The model includes partial structures for the oligo-mannose N-glycans attached to Asn86 (GlcNAc), Asn103 (GlcNAc-GlcNAc), Asn145 (Man-GlcNAc-GlcNAc), Asn291 (GlcNAc), Asn354 (GlcNAc) and Asn374 (GlcNAc) of Clu mutant TL1. The model has excellent stereochemistry with 98.0% of the residues in the favored regions of the Ramachandran plot and no outliers.

The crystal structure of Clu mutant TL1 (Clu-Δ(214–238)) in space group *C*2 was solved by molecular replacement with Molrep 11.4.06 (Vagin and Isupov 2001) using a preliminary model for Clu mutant TL1 in the *P*21 crystal lattice. WinCoot 0.9.4.1 was employed for manual model building (Emsley and Cowtan 2004). The model was refined with Refmac5 5.8.0267 using local non-crystallographic symmetry (NCS) restraints (Murshudov, Skubak et al. 2011). The final refinement was performed with Phenix.refine 1.19.2-4158 (Liebschner, Afonine et al. 2019).

Residues facing solvent channels with disordered side-chains were truncated after C-β. The final model contains two copies of Clu mutant TL1 per asymmetric unit. Residues 23-24, 197-211, 261-279 and 449 in chain A and residues 195-211, 260-279 and 446-449 in chain D, respectively, are missing for lack of discernible electron density, probably because of disorder. The model includes partial structures for the oligo-mannose N-glycans attached to Asn86, Asn103, Asn145, Asn291, Asn354 and Asn374 in chains A and D of TL1. The model has reasonable stereochemistry with 95.2% of the residues in the favored regions of the Ramachandran plot and 0.28% outliers.

### Structure analysis

Coordinates were aligned with Lsqkab 7.1.010 (CCP4i interface 7.1.010) and Lsqman 081126/9.7.9 (Kleywegt and Jones 1994). Molecular drawings and sequence alignment depictions were generated with the programs Pymol 2.2.3 (http://www.pymol.org) and ESPript 3.0 (https://espript.ibcp.fr) (Gouet et al. 1999), respectively.

### Sequence data analysis

An alignment of 242 representative Clu sequences was created with the Consurf server (https://consurf.tau.ac.il/consurf_index.php) (Ashkenazy et al. 2016) using human Clu as input with the default settings. The amino acid frequency of sequences corresponding to residues 204– 238 was analyzed. Sequences for human α-crystallin A and B homologs in jawed vertebrates (taxon ID 7776) were retrieved from Uniprot using BLAST (https://www.uniprot.org/blast), yielding 662 CRYAA/HSPB4 and 444 CRYAB/HSPB5 sequences, respectively, and aligned with Clustalo (https://www.ebi.ac.uk/jdispatcher/msa/clustalo). The N-terminal domains, i.e. the sequences before the α-crystallin domain, which begins at residue 62 in human HSPB4 and residue 65 in human HSPB5, respectively, were cut from the alignments and analyzed. As reference data, the reviewed proteome sequences of *Homo sapiens* and the predicted proteome sequences of *Callorhinchus milii* (Taxonomy ID, 7868), *Latimeria chalumnae* (Taxonomy ID, 7897), *Danio rerio* (Taxonomy ID, 7955), *Xenopus laevis* (Taxonomy ID, 8355), *Gallus gallus* (Taxonomy ID, 9031), *Ornithorhynchus anatinus* (Taxonomy ID, 9258) and *Monodelphis domestica* (Taxonomy ID, 13616) from Uniprot were used.

### Rhodanese aggregation assay

Rhodanese (60 μM) was denatured in 6 M guanidinium-HCl, 5 mM DTT for 1 h at 25 °C and diluted 120-fold into PBS pH 7.2 (Gibco), PBS pH 7.4 (in-house) or 20 mM Na-acetate pH 5.2, 150 mM NaCl, 2 mM CaCl2 (final concentration 0.5 μM) in the absence or presence of Clu constructs (0.5 μM or 1.5 μM). Aggregation was monitored immediately after dilution at 25 °C by measuring turbidity at 320 nm wavelength. The data was fitted using Sigma plot 14.0 software (https://grafiti.com/sigmaplot-detail/) (Exponential Rise to Maximum, Double, 4 Parameter function) to obtained the maximum plateau absorbance.

### Protein aggregation reactions and thioflavin-T (ThT) fluorescence measurements

#### Aβ (M1-42) aggregation

Aβ aggregation assays were performed as previously described (Linse 2020). 500 µg of lyophilized Aβ was resuspended in 700 µl GE buffer pH 8.5 (6 M guanidinium hydrochloride, 20 mM sodium phosphate) and loaded onto a Superdex 75 SEC column (GE Healthcare) equilibrated with aggregation buffer (20mM sodium phosphate, 0.2 mM EDTA, pH 8.5).

Monomeric Aβ was collected in low binding tubes and the concentration was estimated by absorbance at 205 nm. Aβ monomers were diluted to 4 µM in aggregation buffer containing 10 µM ThT in the absence or presence of WT-Clu or Clu mutant proteins. 80 µl of the mix was dispensed per well in a 96 well half-area plate of black polystyrene with a clear bottom (Corning, 3881). Samples were measured in quadruplicates in each plate (technical replicates). ThT signal (excitation 440 nm, emission 480 nm) was measured every 3 min in a CLARIOstar plate reader (BMG Labtech) at 37 °C. The data was fitted using Sigma plot 14.0 software (https://grafiti.com/sigmaplot-detail/) (Sigmoidal, Sigmoid, 3 Parameter function) to obtain the half time for reaching the aggregation plateau.

#### Tau aggregation

For assessing tau aggregation, 80 μl of 10 μM TauRD, 2.5 μM heparin (Merck, H3393), 2 mM MgCl2, 10 μM ThT, PBS 1x pH 7.2 in the presence or absence of 1 μM Clu were dispensed per well in a 96 well half-area plate of black polystyrene with a clear bottom (Corning, 3881).

Samples were measured in quadruplicates in each plate (technical replicates). ThT signal (excitation 440 nm, emission 480 nm, with gain regulation) was measured every 2 min in a SPARK multimode microplate reader (TECAN) at 37 °C under constant shaking (50 s linear shaking: amplitude 4.5 mm, frequency 420 rpm - 50 s orbital shaking: amplitude 1.5 mm, frequency 360 rpm). The data was fitted using Sigma plot 14.0 software (Sigmoidal, Sigmoid, 3 Parameter function) to obtain the half time for reaching the aggregation plateau. A detailed protocol is available here: dx.doi.org/10.17504/protocols.io.dm6gp3nw8vzp/v1.

#### α-Synuclein aggregation

For assessing α-synuclein aggregation, 80 μl of 200 μM α-synuclein, 0.05% NaN3, 10 μM ThT, 150 mM KCl, 50 mM Tris-HCl pH 7.6 in the presence or absence of Clu at 0.04 μM were dispensed per well in a 96 well half-area plate of black polystyrene with a clear bottom (Corning, 3881). Samples were measured in quadruplicates in each plate (technical replicates). ThT signal (excitation 440 nm, emission 480 nm, with gain regulation) was measured every 10 min in a SPARK multimode microplate reader (TECAN) at 37 °C under constant shaking (120 s linear shaking: amplitude 1.5 mm, frequency 1080 rpm - 120 s orbital shaking: amplitude 1 mm, frequency 510 rpm). The data was fitted using Sigma plot 14.0 software (https://grafiti.com/sigmaplot-detail/) (Sigmoidal, Sigmoid, 3 Parameter function) to obtain the half time for reaching the aggregation plateau. A detailed protocol is available here: dx.doi.org/10.17504/protocols.io.8epv5x87dg1b/v1.

### Protein labeling

Clusterin was labeled with Alexa488 N-hydroxysuccinimide ester (Thermo Fisher Scientific). Before the labeling reaction, the buffer was exchanged with 0.1 M sodium bicarbonate buffer pH 8.3 (N-terminal labeling buffer) using a Nap5 column and labeling was subsequently performed at a 4-fold molar excess of Alexa 488 for 1.5 h at RT. Free dye was removed using a Nap5 column, pre-equilibrated with PBS buffer. The labeling efficiency was measured by nanodrop and was typically about 70–180% (note that Clu contains 2 N-termini). A detailed protocol is available here: dx.doi.org/10.17504/protocols.io.rm7vzxpbrgx1/v1.

### Cellular uptake assays

5µg ml^−1^ of clusterin-A488 in 400 µl medium (200 µl of fresh media) was added to 250,000 iNeurons cultured in a well of a 12-well plate. After 1 h, cells were placed on ice, washed with PBS and collected with Accutase (Stem Cell technologies). Cells were washed once with PBS, fixed with 4% PFA/PBS for 10 min, washed with PBS, resuspended in 160 µl of PBS and stored at 4 °C until analysis. Cells were analyzed at the MPIB Imaging Facility with an Attune NxT flow cytometer (Thermo Fisher Scientific). Right before measuring, 50 µl of Trypan blue solution 0.4% (Thermo Fisher Scientific) were added to each sample to quench the A488 fluorescence outside the cells. Uptake was recorded during the linear increase of the signal, i.e. before degradation became apparent. To measure the A488 signal, cells were excited with 488 nm laser light and fluorescence was determined using the 530/30 filter. For each sample at least 10,000 cells were analyzed (average analyzed cells: ∼98,000). Data processing was performed using MatLabR2021b (program code at https://github.com/csitron/MATLAB-Programs-for-Flow-Cytometry). Cells were gated by size using forward scatter (FSC-H) and A488 mean intensity normalized by FSC-H of each mutant was normalized by their own labeling efficiency.

### Immunofluorescence microscopy

100,000 iNeurons were cultured in a well of a 24-well plate on 13 mm coverslips. Cells were washed with PBS, fixed with 4% PFA/PBS for 10 min, washed with PBS and permeabilized with 0.1% Triton-X100/PBS for 5 min. Blocking solution (8% BSA/PBS) was added for 1 h.

Coverslips were transferred to a humid chamber and incubated overnight with the primary antibody diluted in 1% BSA/PBS (anti-MAP2 antibody (AB554, Merck), 1/500 dilution; anti-β- 3-tubulin (MA1-19187, Thermo Fisher Scientific), 1/100 dilution). Cells were then washed with PBS, incubated with the respective secondary antibody, goat anti-chicken IgY (H+L), Alexa Fluor 647 (A-21449, Thermo Fisher Scientific) or F(ab’)2-goat anti-mouse IgG (H+L), Alexa Fluor Plus 647 (A48289, Thermo Fisher Scientific, 1/500 dilution), diluted in 1% BSA/PBS for 1 h, washed with PBS and stained with NucBlue fixed cell ReadyProbes reagent (Thermo Fisher Scientific). Coverslips were mounted with Dako fluorescence mounting medium (Agilent). The confocal imaging was performed at the MPIB Imaging Facility, on a LEICA TCS SP8 AOBS confocal laser scanning microscope (Wetzlar, Germany) equipped with a LEICA HCX PL APO 63x/NA1.4 oil immersion objective. Images were analyzed with Image J (https://imagej.net/ij/). A detailed protocol is available here: dx.doi.org/10.17504/protocols.io.e6nvwd7n7lmk/v1, except that blocking buffer was 8% BSA/PBS and antibody buffer 1% BSA/PBS.

### Clu–VLDLR-ed binding assay

WT-Clu in presence and absence of VLDLR-ed, all at 5 μM, were incubated with 25 μl CaptureSelect C-tag affinity resin (Thermo Fisher Scientific) in C-tag wash buffer (20 mM Tris-HCl pH 7.4, 100 mM NaCl and 2 mM CaCl2) for 2 h at 25 °C, followed by transfer into spin columns (Mo Bi Tec). Subsequently, the gel bed was washed four times with 100 μl C-tag wash buffer. Bound protein was eluted with three times 50 μl C-tag Elution buffer (20 mM Tris-HCl pH 7.0, 2 M MgCl2 and 2 mM CaCl2). Protein association was analyzed by SDS-PAGE and immunoblotting against Clu α-chain and C-tag.

### Solid phase binding assay

The assay was performed as previously described (Leeb, Eresheim and Nimpf 2014). The 96-well plate (Nunc-immuno MicroWell 96 well solid plate, Merck) was coated with 100 µl of TBS-C (Tris-buffered saline pH 7.4, 2 mM CaCl2) containing 10 µg ml^−1^ VLDLR-ed overnight at 4 °C. The plate was washed once with TBS-C and incubated for 2 h at RT with TBS-C blocking buffer (2% BSA, 0.05% Tween). The plate was then incubated with different concentrations of the ligands Clu or RAP in blocking solution during 1 h at RT. For the competition assay, the plate was incubated with 100 nM of Clu in the presence of increasing concentrations of RAP. The plate was washed three times with blocking solution. Anti-clusterin (sc-5289, Santa Cruz Biotechnologies; dilution 1/100) or anti-RAP (sc-515625, Santa Cruz Biotechnologies; dilution 1/100) antibodies were added in TBS-C blocking solution and incubated during 1 h at RT. The plate was washed three times with TBS-C blocking solution and incubated with goat-anti mouse IgG-HRP (A4416, Merck; dilution 1/10,000) added in TBS-C blocking solution and incubated during 1 h at RT. The plate was washed three times with TBS-C blocking solution, developed by adding 100 µl per well of the HRP substrate 1-Step Ultra TMB ELISA Substrate Solution (Thermo Fischer Scientific) and the reaction was quenched with 100 µl per well of 2 M sulfuric acid. Absorbance at 450 nm was measured in a SPARK multimode microplate reader (TECAN). Binding to wells coated with BSA was used to estimate the background signal for each sample and no-ligand well signal was subtracted from the rest of the samples. The dissociation constants (*K*D values) were obtained by fitting the data using GraphPad Prism 10 software (www.graphpad.com) (Binding – saturation, one site – specific binding). A detailed protocol is available here: dx.doi.org/10.17504/protocols.io.yxmvm36kol3p/v1.

### Formation and isolation of protein-phospholipid particles

1,2-dimyristoyl-sn-glycero-3-phosphocholine (DMPC, Avanti polar lipids, Sigma) powder was dissolved in 3:1 chloroform/methanol at 25 mg ml^−1^ or 5 mg ml^−1^ and stored at –70 °C as stock solution. For the formation of Clu–phospholipid particles, the required amount of DMPC was transferred to a glass vial and solvent was removed by evaporation through a constant stream of nitrogen gas. The dried film was resuspended in 1x PBS pH 7.2 (Gibco), vortexed and sonicated in a Bioruptor sonication bath (Diagenode) (25 cycles of 5 s on – 5 s off). 20 µM Clu, Hsc70, ApoA1 (CYT-661, Prospec, Hölzel Diagnostik), the respective Clu-GFP construct or 6 µM ApoE (CYT-874, Prospec, Hölzel Diagnostik) were then mixed (1:1, v/v) at the indicated ratios with DMPC in a total volume of 100 µl. For DMPC–Clu lipoprotein complex isolation by SEC, 20 µM Clu or Clu-A488 were mixed with 20 or 10 mM DMPC. The sample was incubated through 3 cycles of 18 °C for 15 min and 30 °C for 15 min, as described (Yeh, Wang et al. 2016). The formation of the Clu–phospholipid particles was analyzed by native PAGE. Coomassie-blue protein staining was performed with InstantBlue (Abcam) or with 0.1% (w/v) Serva Blue R in 10% acetic acid and 50% ethanol followed by de-staining with 10% acetic acid and 10% ethanol. The lipoprotein complex band formed in presence of Clu mutant proteins was quantified by densitometry with Image J. The detection of lipids after native PAGE was performed by incubating the gel overnight with freshly prepared 0.4% Sudan black B (Merck) in 16.7% acetone, 12.5% acetic acid solution and de-staining with 20% acetone, 15% acetic acid. When Clu-A488 or the GFP constructs were used, the fluorescence signal of the native PAGE gel was analyzed using a Typhoon 5 imager (Cytiva).

When Clu–phospholipid particles were isolated by SEC, after lipidation, the sample was briefly centrifuged to remove large particles and the supernatant was loaded onto a Superose 6 column equilibrated with 1x PBS or TBS-C (for chymotrypsin digestion). After analysis by native PAGE, fractions containing the Clu–DMPC nanodisc particles were concentrated by ultrafiltration using Vivaspin MWCO 30,000 or 10,000 (GE Healthcare) centrifugal concentrators. Protein concentration was determined by absorbance at 280 nm or with the Bradford assay, and lipid content was analyzed by mass spectrometry (see below) and with the Phospholipid Assay Kit (MAK122, Merck). A detailed protocol is available here: dx.doi.org/10.17504/protocols.io.bp2l6x59zlqe/v1.

### Negative stain transmission electron microscopy

For negative stain analysis, continuous carbon grids (Quantfoil) were glow discharged using a plasma cleaner (PDC-3XG, Harrick) for 30 s. Grids were incubated for 1 min with 4 μl of the fractions of the Clu–DMPC band from SEC (in PBS buffer) or the GFP fusion protein αTL-H7 lipoprotein complex preparation, blotted and stained with 2% uranyl acetate solution (Electron Microscopy Sciences), dried and imaged at the MPIB Cryo-EM Core Facility, on a Titan Halo (FEI) transmission electron microscope using SerialEM v. 4.1.6.

### Mass spectrometry lipidomics

Mass spectrometry lipidomics was performed at the MPIB Mass Spectrometry Core Facility.

#### Lipid extraction

Lipids (from proteins or DMPC standards) were extracted using a methyl tert-butyl ether (MTBE)-based extraction method. A volume of 200 µl of cold methanol and 800 µl of cold MTBE were added, and the samples were vortexed. After adding 200 µl of water, a phase separation appeared. The extraction mixture was centrifuged at 10,000 x g for 10 min at 4 °C to separate the organic and aqueous phases. The upper organic phase was collected and dried by vacuum centrifugation.

#### LC-MS/MS data acquisition and analysis

Data were recorded using a QExactive HF mass spectrometer (Thermo Scientific) coupled to a Vanquish Flex HPLC system (Thermo Fisher Scientific). The lipid extracts and DMPC standard extracts were reconstituted in 40 µl or 20 µl of acetonitrile/isopropanol/water in a 65:30:5 ratio (v/v/v), respectively. 1 µl of the samples or 10 µl of the standards were injected and separated on a C8 column (Luna 3u, 2 x 100 mm, 3.0 µm, Phenomenex) at a flow rate of 150 µl min^−1^. Mobile phases A and B consisted of acetonitrile 60:40% (v/v) and isopropanol 90:10% (v/v), both buffered with 0.1% formic acid and 10 mM ammonium formate. Buffer B was maintained at 30% for 1 min, then increased to 61% within 7 min, further increased to 71% within 6 min, and finally increased to 99% within 4 min. The percentage of buffer B was held at 99% for 4.5 min. The column was then re-equilibrated to 30% B for 2.5 min.

The mass spectrometer operated in positive mode with data-dependent MS1 scans from 160 to 1600 m/z at a resolution of 120,000. Conditions for the HESI source were as followed: Sheath gas (N2) flow rate was set to 47 (arbitrary units), auxiliary gas flow rate was set to 10 (arbitrary units), and sweep gas flow rate was set to 2 (arbitrary units). The spray voltage was maintained at 3.20 kV, and the capillary temperature was set to 250 °C and the temperature of the auxiliary gas heater was set to 380 °C. Up to 5 of the top precursors were selected and fragmented using higher energy collisional dissociation (stepped-HCD with normalized collision energies of 20, 40, and 60). The MS2 spectra were recorded at a resolution of 30,000. The AGC target for MS1 and MS2 scans was set to 3E6 and 1E5, respectively, within a maximum injection time of 200 ms for MS and 50 ms for MS2 scans.

Peak areas corresponding to DMPC were identified based on MS1 high-resolution mass and retention time (previously identified using unlabeled standards) using the software "Skyline," version 23.1.0.455 (https://skyline.ms/). Exact amounts were calculated based on a previously recorded calibration curve.

### Optiprep gradient

Rhodanese aggregation reactions in the presence of Clu or Clu–DMPC at molar ratio Clu/D-Rho 1:3 were centrifuged at 20,000 x g for 15 min at 4 °C to remove large aggregates. 60 µl of the supernatants were mixed with Optiprep (Merck) to a final concentration of 36% in PBS and final volume of 1 ml. The sample was placed in a 3.5 ml thick wall polycarbonate tube (349622, Beckman Coulter). 1 ml of 24% Optiprep diluted in PBS was placed carefully on top and 1 ml of PBS was added as top layer. The gradient was centrifuged at 54,000 rpm for 3 h at 4 °C using a SW55 Ti rotor (Beckman Coulter). After centrifugation, six fractions of 0.5 ml were manually collected, diluted with 0.5 ml PBS and 40 µl of 2% Na-deoxycholate were added. After 15 min incubation on ice, 100 µl of 100% trichloroacetic acid were added, followed by 1 h incubation on ice. The samples were then centrifuged at 20,000 x g for 30 min at 4 °C. The pellets were washed with 500 μl ice-cold acetone, sonicated in a Bioruptor sonication bath (Diagenode) (2 cycles of 30 s on – 30 s off) and centrifuged at 20,000 x g for 10 min at 4 °C. The pellets were air dried and 30 μl 300 mM Tris-HCl pH 8.8 were added and incubated 5 min on ice. 30 μl NuPAGE LDS Sample Buffer (4X) buffer (Thermo Fisher Scientific) containing 100 mM DTT were added and the mixture boiled for 10 min, followed by SDS-PAGE analysis and immunoblotting against Clu and rhodanese.

### Limited proteolysis with chymotrypsin

Free WT-Clu or TL4 mutant or their respective Clu–DMPC lipoprotein complex preparations at 10 μM Clu content were incubated with a series of bovine chymotrypsin (Merck) concentrations (0, 1, 2, 5, 10, 20, 50 nM) in TBS buffer containing 1 mM CaCl2 for 30 min at 25 °C. The protease reactions were stopped on ice by addition of PMSF (final concentration 10 mM), followed by SDS-PAGE and native PAGE analysis. Selective bands were excised and the protein content recovered, digested with trypsin and analyzed by mass spectrometry.

### Mass spectrometry proteomics

Mass spectrometry proteomics was performed at the MPIB Mass Spectrometry Core Facility.

#### Sample preparation for proteomics

The gel pieces were thoroughly washed multiple times with 150 µl of a destaining buffer (containing 25 mM ammonium bicarbonate and 50% ethanol) and then dehydrated with 150 µl of pure ethanol. After ethanol removal, the gel pieces were dried using vacuum centrifugation. Then, 50 µl of a digestion buffer (25 mM Tris-HCl, 10% acetonitrile, 10 ng µl^−1^ trypsin) was added. The mixture was cooled on ice for 20 min, followed by the addition of 50 µl of 25 mM ammonium bicarbonate buffer. The gel pieces were incubated overnight at 37 °C. The peptides in the supernatant were collected, and additional peptides were extracted by repeated incubation at 25 °C in 100 µl of an extraction buffer (3% trifluoroacetic acid (TFA), 30% acetonitrile), followed by centrifugation and collection of the supernatants. Finally, the gel pieces were dehydrated by incubation at 25 °C in 100 µl of pure acetonitrile, and the supernatant was combined with those from the previous steps. Acetonitrile was removed by vacuum centrifugation, and 70 µl of a solution containing 2 M Tris-HCl, 10 mM TCEP, and 40 mM chloroacetamide (CAA) was added. After a 30 min incubation at 37 °C, the peptides were acidified to 1% TFA.

#### LC-MS/MS data acquisition

Desalted peptides were separated on a 30 cm column (75 µm inner diameter) packed with ReproSil-Pur C18-AQ 1.9-µm beads (Dr. Maisch GmbH) at a flow rate of 300 nl min^−1^ using a Thermo Easy-nLC 1200 system at 60 °C. Peptides were directly introduced into the Exploris 480 mass spectrometer via a nano-electrospray interface. The LC-MS gradient used buffer A (0.1% formic acid) and buffer B (80% acetonitrile, 0.1% formic acid): buffer B was increased from 5% to 30% over 30 min, then to 65% in 5 min, and finally to 95% over the next 5 min, maintaining 95% for an additional 5 min. The mass spectrometer was operated in data-dependent mode with survey scans from 300 to 1650 m/z (resolution of 60,000 at m/z 200). Up to 15 top precursors were selected and fragmented using higher-energy collisional dissociation (HCD) with a normalized collision energy of 28. MS2 spectra were recorded at a resolution of 15,000 (at m/z 200). AGC targets for MS and MS2 scans were set to 3E6 and 1E5, with maximum injection times of 25 ms and 28 ms, respectively. Dynamic exclusion was set to 30 s.

#### Mass spectrometry data analysis

Raw data were processed using the MaxQuant computational platform (version 2.2.0.0, https://www.maxquant.org/) (Cox and Mann 2008) with standard settings applied. Briefly, the peak list was searched against the UniProt sequences of human proteins (SwissProt and TrEMBL) with an allowed precursor mass deviation of 4.5 ppm and an allowed fragment mass deviation of 20 ppm. Cysteine carbamidomethylation, methionine oxidation, and N-terminal acetylation were set as variable modifications. Trypsin/P was set as the protease, and the digestion mode was set to "semi-specific".

### Statistical analysis

Statistical analysis was performed with GraphPrism10 (www.graphpad.com). Sample size (*n*) given in figure legends describe measurements taken from distinct, independent samples. One-way ANOVA with Dunnett’s post hoc test was used for multiple comparisons. Exact p-values indicated in Source data.

## Acknowledgements

We thank Theresa F. Schaller and David Balchin for their contributions to the early phase of the Clu structural project; Tim Klaubert, Lavina Dinata, George-Valentin Datcu, Justin Samiee, Aysegül Eylül Agaciklar and Devika Rege for their help in the preparation and characterization of Clu mutants and GFP fusion proteins; Nadine Wischnewski and Silvia Gärtner for purification of Aβ(M1-42) and Hsc70, and RAP proteins, respectively; Roman Körner and Albert Ries for help with mass spectrometry experiments; Cole Sitron for sharing the flow cytometry analysis code and for helpful discussions; Dominik Paquet and his team for kindly sharing their expertise on iPSCs culture and differentiation protocols. We acknowledge the European Synchrotron Radiation Facility for provision of macromolecular crystallography infrastructure, and we would like to thank Daniele de Sanctis for assistance in using beamline ID23-1. We thank the staff of the Max Planck Institute of Biochemistry (MPIB) Core, Cryo-EM and Crystallization facilities, especially Judith Scholz, for generating the HEK293-EBNA cell lines and expressing the Clu and VLDLR-ed constructs; Martin Spitaler, Markus Oster, and Giovanni Cardone from the MPIB Imaging facility for support with flow cytometry, imaging and image processing; Barbara Steigenberger and her team from the MPIB Mass Spectrometry (MS) core facility for mass spectrometry lipidomics and proteomics analysis. This research was funded in part by the Deutsche Forschungsgemeinschaft (DFG, German Research Foundation) under Germany’s Excellence Strategy within the framework of the Munich Cluster for Systems Neurology (EXC 2145 SyNergy—ID 390857198) and by the Aligning Science Across Parkinson’s initiative [ASAP-000282] through the Michael J. Fox Foundation for Parkinson’s Research (MJFF). For the purpose of open access, the authors have applied a CC BY public copyright license to all Author Accepted Manuscripts arising from this submission.

## Author contributions

AB determined and analyzed the clusterin structure and designed the mutant constructs. PY designed, performed and analyzed the biochemical experiments with support from AB and SP. AIC performed H/DX experiments and MS analysis. CM performed the negative stain EM. AB, PY and FUH designed the project and wrote the manuscript with input from the other coauthors.

## Additional information

Correspondence and requests for materials should be addressed to Andreas Bracher, F. Ulrich Hartl or Patricia Yuste-Checa.

## Data availability

The DOI associated with the diffraction data is 10.15151/ESRF-ES-541098252. The coordinates and structure factors reported in this manuscript have been deposited in the Worldwide Protein Data Bank with accession codes 7ZET and 7ZEU. The mass spectrometry data have been deposited to the ProteomeXchange Consortium via the PRIDE partner repository (https://www.ebi.ac.uk/pride/login) with the dataset identifiers PXD056940 (Username, password: FX14UNyJ1WjX) and PXD057022 (Username, password: pR8ZTxHvceFh). MatLabR2021b code for flow cytometry data processing was deposited (https://github.com/csitron/MATLAB-Programs-for-Flow-Cytometry). Detailed information and RRID of chemicals, recombinant proteins, antibodies, commercial assays, recombinant DNA, protocols and cell lines are listed in Supplementary Table 2 (Key resource table).

