## Supplementary Information for "Hydrophobic tails enable diverse functions of the extracellular chaperone clusterin"

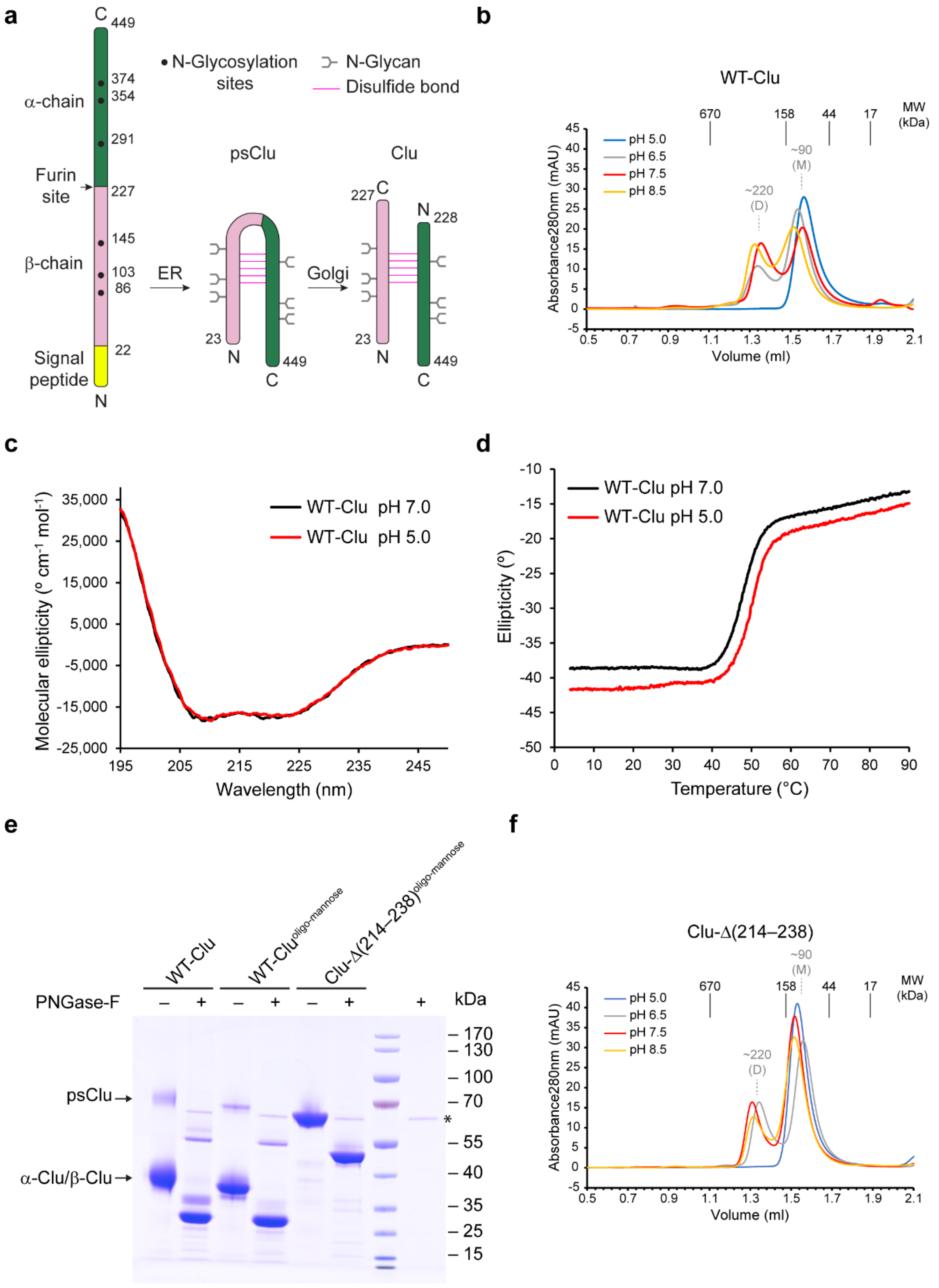

a
C 449
374
354
291
α-chain
Furin site
227
145
β-chain
103
86
22
Signal peptide
N
N-Glycosylation sites
N-Glycan
Disulfide bond
ER
psClu
Golgi
Clu
b
WT-Clu
pH 5.0
pH 6.5
pH 7.5
pH 8.5
670
158
44
17
MW (kDa)
~90 (M)
~220 (D)
Absorbance280nm (mAU)
Volume (ml)
c
WT-Clu pH 7.0
WT-Clu pH 5.0
Molecular ellipticity (° cm-1 mol-1)
Wavelength (nm)
d
WT-Clu pH 7.0
WT-Clu pH 5.0
Ellipticity (°)
Temperature (°C)
e
WT-Clu
WT-Clu oligo-mannose
Clu-Δ(214–238) oligo-mannose
PNGase-F
kDa
psClu
α-Clu/β-Clu
170
130
100
70
55
40
35
25
15
*
f
Clu-Δ(214–238)
pH 5.0
pH 6.5
pH 7.5
pH 8.5
~90 (M)
~220 (D)
MW (kDa)
Absorbance280nm (mAU)
Volume (ml)

**Supplementary Figure 1: Biochemical characterization of recombinant clusterin proteins.** (**a**) Biogenesis of secreted Clu. The Clu precursor is thought to be co-translationally inserted into the endoplasmic reticulum (ER) lumen followed by signal peptide (yellow) removal. In the ER, six N-glycans are attached and five anti-parallel disulfide bonds are introduced (Choi-Miura et al. 1992). During passage through the Golgi network, the N-glycans are processed and a furin-like protease cleaves Clu into β- and α-chains (pink and dark green). N-glycans and disulfide bonds are indicated by forks (gray) and horizontal lines (magenta), respectively. N-glycosylation sites, furin site and chain termini are indicated. (**b**) pH-dependent oligomerization of WT-Clu. WT-Clu at 10 μM concentration was incubated on ice with buffer containing 100 mM NaCl and 1 mM EDTA and either 20 mM Na-acetate pH 5.0, 20 mM MES-NaOH pH 6.5, 20 mM HEPES-NaOH pH 7.5 or Tris-HCl pH 8.5 overnight or longer (see Methods for details). The samples were subsequently analyzed by SEC on a Superdex-200 Increase 3.2/300 column at room temperature (RT). Absorbance traces at 280 nm wavelength are shown. The retention volume of molecular weight standards with their mass values in kDa is indicated in black. The apparent molecular weight of Clu is indicated in grey (M: monomers; D: dimers). Note that the apparent size of Clu observed by SEC is larger than expected due to its elongated shape. Representative results are shown (*n* =3 independent experiments). (**c**) CD spectra of WT-Clu at pH 5.0 and pH 7.0. CD spectra were recorded at 0.1 mg $ml^{-1}$ protein concentration at 20 °C in 50 mM K-phosphate pH 5.0 (red) or pH 7.0 (black). Averages of 3 independently prepared samples are shown. Molecular ellipticities were calculated. (**d**) CD melting curves of WT-Clu at pH 5.0 and pH 7.0. The CD signal at 222 nm wavelength was recorded during slow heating (60 °C $h^{-1}$) of 5 μM of WT-Clu in 50 mM K-phosphate pH 5.0 (red) or pH 7.0 (black). Representative curves are shown (n = 3 experiments). The averages of the melting temperatures are 47.6 and 49.6 °C for pH 7.0 and 5.0, respectively. (**e**) SDS-PAGE analysis of Clu preparations. A Coomassie blue-stained gel is shown. WT-Clu with natural N-glycans and the oligo-mannose forms of WT-Clu and the Clu-Δ(214–238) mutant produced in presence of the α-mannosidase-I inhibitor kifunensine are shown before (respective left lane) and after (respective right lane) treatment with the glycosidase PNGase-F (GST-PNGase-F), which removes N-glycans and replaces the attachment site Asn with Asp. The bands of the α- and β-chains of WT-Clu overlap at ~40 kDa. Residual uncleaved Clu precursor (psClu) is indicated. The WT-Clu bands before PNGase-F cleavage are fuzzy because of glycan heterogeneity. On the two rightmost lanes, molecular weight markers and GST-PNGase-F (asterisk) were analyzed. (**f**) pH-dependent oligomerization of the Clu-Δ(214–238) mutant. The experimental setup was identical to that for WT-Clu (panelb). Representative results are shown (*n* =3 independent experiments).

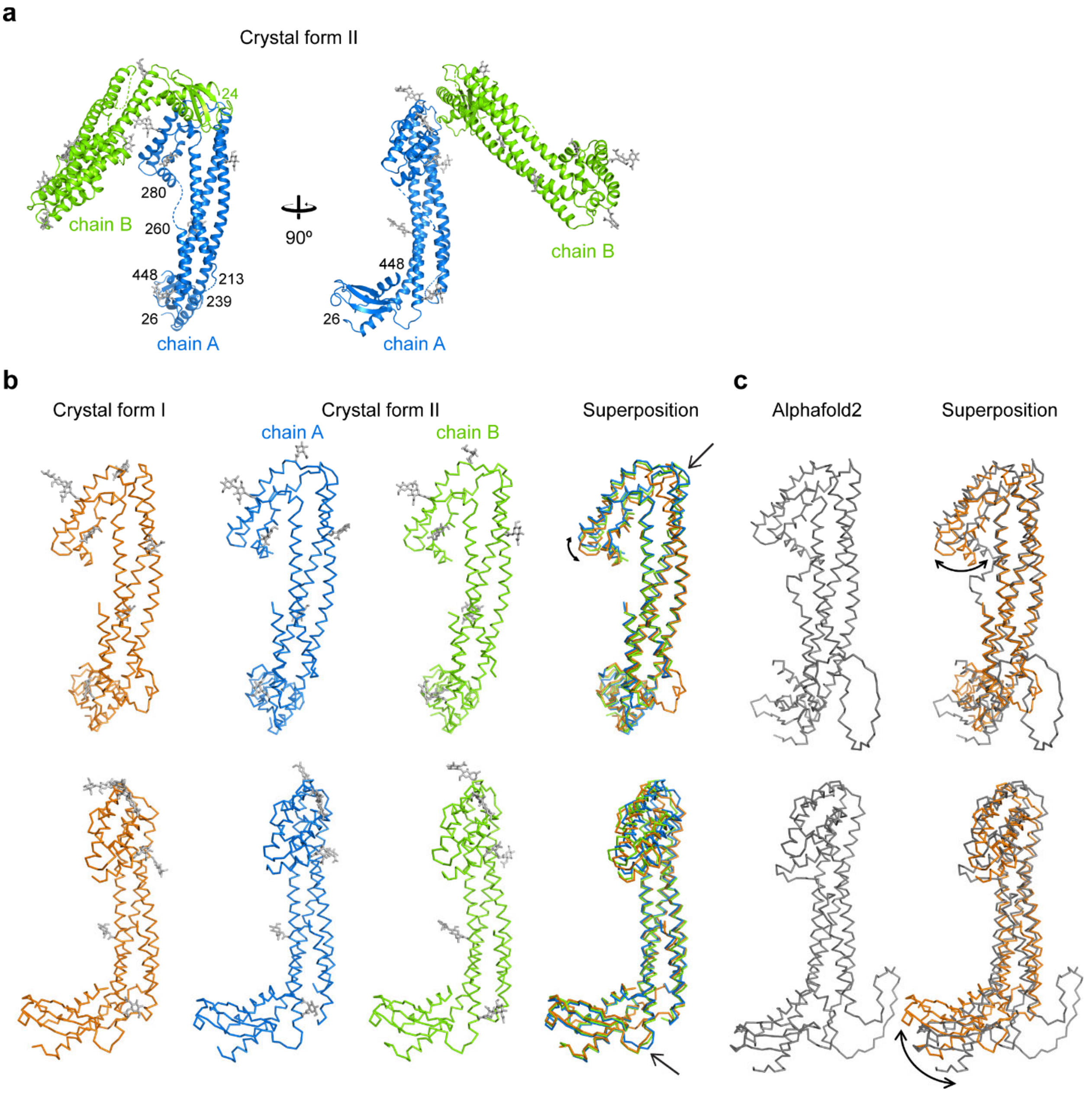

**Supplementary Figure 2: Comparison of clusterin crystal structures.** (**a**) The asymmetric unit of crystal form II. Perpendicular views are shown. Copies A and B of Clu-Δ(214–238) mutant protein are shown as ribbons in blue and green, respectively. First and last ordered residue and gaps in the model are indicated. The ordered parts of N-glycans are shown in stick representation in gray. (**b**) Comparison of crystallographically independent copies of Clu. The three independent copies of Clu-Δ(214–238) mutant protein in crystal forms I and II are shown in orange, blue and green, respectively, on the left. On the right, a superposition is shown. The peptides are shown as Cα traces and N-glycans as sticks. Arrows indicate regions of local divergence. Two perpendicular views are shown. (**c**) Comparison of the experimental Clu structure with the Alphafold2 model. The Alphafold2 model for human Clu (https://www.alphafold.ebi.ac.uk/entry/P10909) is shown in gray on the left and a superposition with Clu-Δ(214–238) mutant protein in crystal form I on the right. The disulfide domain and the α/β roll-like domain exhibit noticeable reorientations in the Alphafold2 model (indicated by curved arrows).

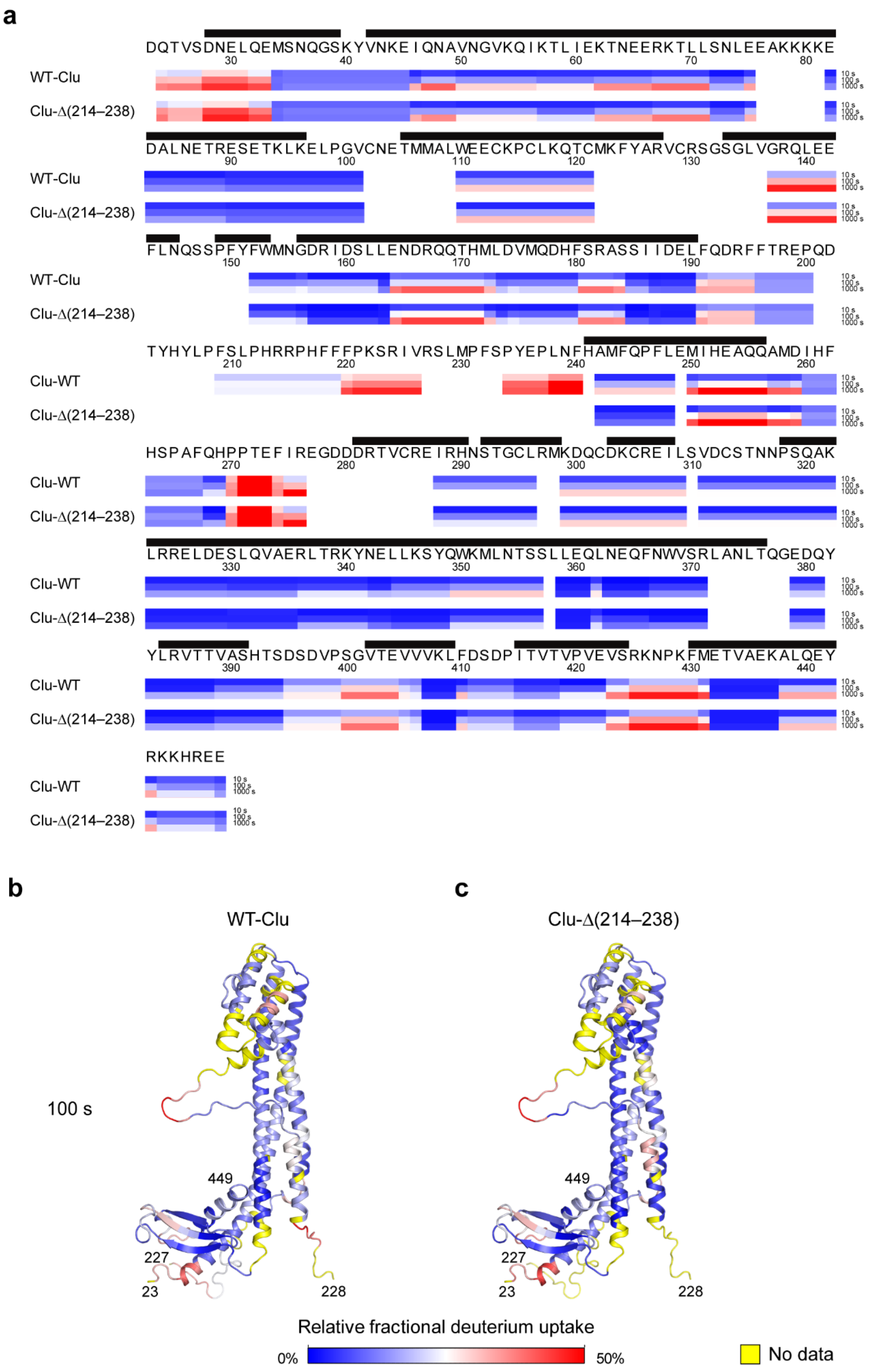
a
DQTVSDNELQEMSNQGSKYVNKEIQNAVNGVKQIKTLIEKTNEERKTLLSNLEEAKKKKE
30 40 50 60 70 80
WT-Clu
Clu-Δ(214–238)
DALNETRESETKLKELPGVCNETMMALWEECKPCLKQTCMKFYARVCRSGSGLVGRQLEE
90 100 110 120 130 140
WT-Clu
Clu-Δ(214–238)
FLNQSSPFYFWMNGDRIDSLLENDRQQTHMLDVMQDHFSRASSIIDELFQDRFFTREPQD
150 160 170 180 190 200
WT-Clu
Clu-Δ(214–238)
TYHYLPFSLPHRRPHFFFPKSRIVRSLMPFSPYEPLNFHAMFQPFLEMIHEAQQAMDIHF
210 220 230 240 250 260
Clu-WT
Clu-Δ(214–238)
HSPAFQHPPTEFIREGDDDRTVCREIRHNSTGCLRMKDQCDKCREILSVDCSTNNPSQAK
270 280 290 300 310 320
Clu-WT
Clu-Δ(214–238)
LRRELDESLQVAERLTRKYNELLKSYQWKMLNTSSLLEQLNEQFNWVSRLANLTQGEDQY
330 340 350 360 370 380
Clu-WT
Clu-Δ(214–238)
YLRVTTVASHTSDSDVPSGVTEVVVKLFDSDPITVTVPVEVSRKNPKFMETVAEKALQEY
390 400 410 420 430 440
Clu-WT
Clu-Δ(214–238)
RKKHREE
Clu-WT
Clu-Δ(214–238)
10 s
100 s
1000 s
b
c
WT-Clu
Clu-Δ(214–238)
100 s
449
227
23
228
Relative fractional deuterium uptake
0%
50%
No data

**Supplementary Figure 3: Hydrogen-deuterium exchange analysis of clusterin.** (**a**) Peptide-level H/DX kinetics for WT-Clu (upper rows) and Clu-Δ(214–238) mutant (lower rows) represented as heat maps. WT-Clu or Clu-Δ(214–238) mutant at pH 5.0 were exposed to deuterated buffer for 10 s (top), 100 s (middle) and 1000 s (bottom), respectively. After quenching exchange at pH 2.6 and 0 ºC, the protein was digested with pepsin and deuterium incorporation into backbone amides of fragment peptides determined by MS. For the analysis, the data from three independent experiments were used. The peptides considered for analysis were identified at least twice. The blue-white-red color gradient represents increasing fractional deuterium incorporation from 0 to 50 %. Secondary structure (black bars), sequence and residue numbers of WT-Clu are indicated. Slow and rapid exchange kinetics correlated well with presence of secondary structure and disorder, respectively. For unknown reasons, peptides 196–200, 209–219 and 260–269, predicted to be disordered, exhibited near constant intermediate deuterium levels at all time points. (**b, c**) Mapping of the hydrogen-deuterium exchange data onto the Clu structure model. The fractional deuterium incorporation into peptide segments during 100 s in WT-Clu (b) and Clu-Δ(214–238) (c) was mapped onto the structure model. The same views as in the right panel of Figure 1b are shown. The blue-white-red color gradient (see scale bar) represents increasing fractional deuterium incorporation from 0 to 50 %. Yellow coloring indicates missing data coverage. N- and C- terminal residues are indicated. Residues 28–33 in helix α1 and residues 137–142 in helix α4 exhibited increased H/D exchange in both samples, consistent with enhanced structural plasticity in solution compared to the crystal lattice.

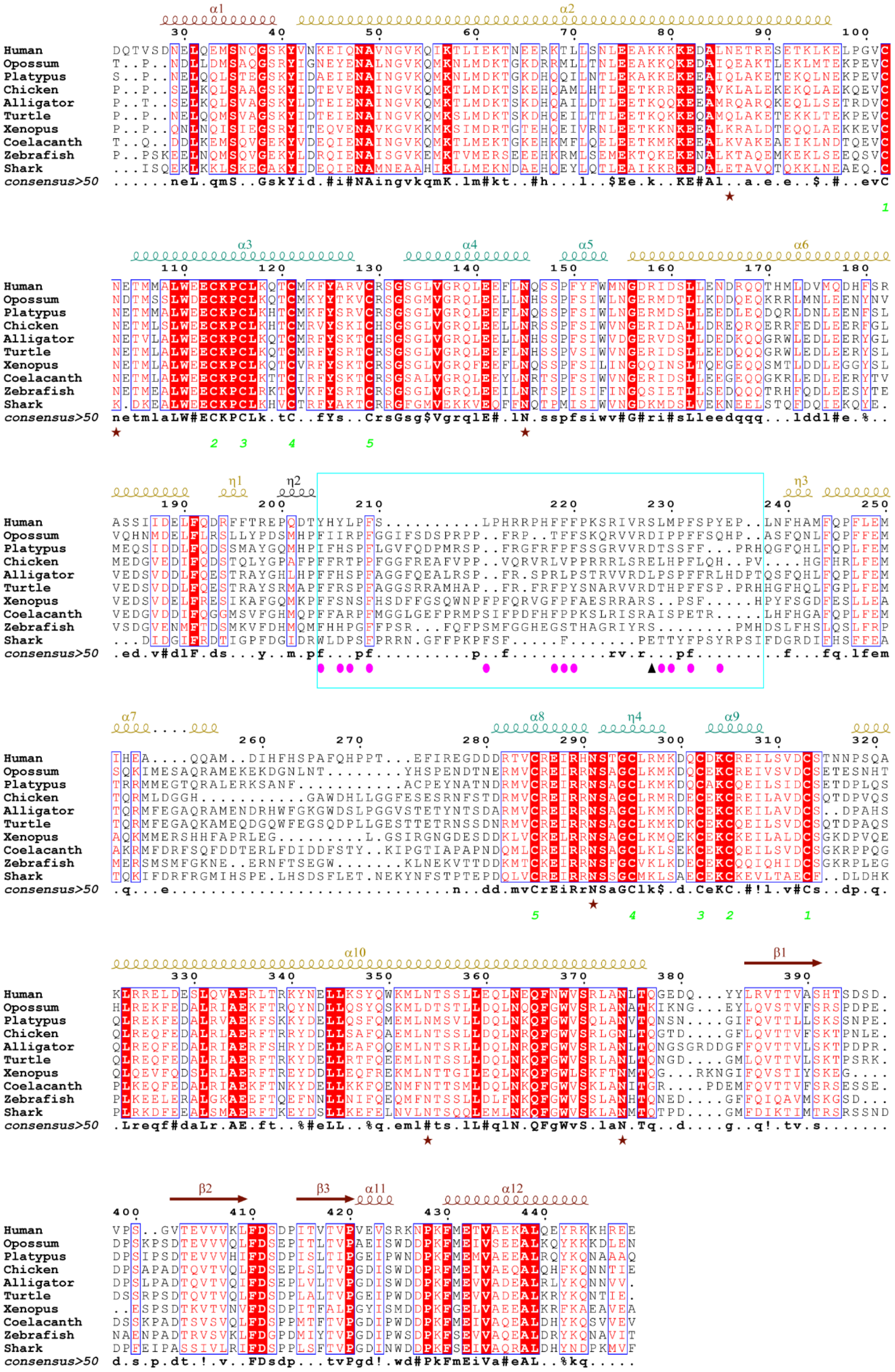

α1
α2
α3
α4
α5
α6
η1
η2
η3
α7
α8
η4
α9
α10
β1
β2
β3
α11
α12
Human
Opossum
Platypus
Chicken
Alligator
Turtle
Xenopus
Coelacanth
Zebrafish
Shark
consensus>50

**Supplementary Figure 4: Alignment of representative clusterin sequences.** Amino acid sequences of a representative set of Clu homologs were aligned using the EBI Clustal-Ω server (https://www.ebi.ac.uk/Tools/msa/clustalo/). Secondary structure elements for human Clu are indicated above the sequences. The Clu domain structure is indicated by dark red, gold and teal coloring of secondary structure elements in the α/β roll-like, coiled-coil and disulfide domain, respectively. Similar residues are shown in red and identical residues in white on a red background. Blue frames indicate homologous regions. The consensus sequence is shown at the bottom (Upper case, conserved; lower case, conserved in more than 50% of the sequences; symbol explanations: #, D/N/E/Q, $, L/M; % F/Y; !, I/V). The ER-targeting signal sequences are not shown. Asterisks in brown below the sequence indicate attachment sites for N-glycans in human Clu. Disulfide bonds 1–5 are represented by green numbers. Residues mapping to the disordered tails are indicated by a frame (cyan). The furin cleavage site is indicated by an arrowhead (black). Hydrophobic and aromatic residues in the tail regions that were mutated in this study are indicated by ovals in pink. Uniprot accession codes for the sequences are: P10909, *Homo sapiens* (human); F6UX07, *Monodelphis domestica* (opossum); F6XFZ6, *Ornithorhynchus anatinus* (platypus); A0A1D5PN61, *Gallus gallus* (chicken); A0A1U7SNW7, *Alligator sinensis* (chinese alligator); XP_005284162.1, *Chrysemys picta bellii* (painted turtle); Q6DIX4, *Xenopus tropicalis* (western clawed frog); H3AKI1, *Latimeria chalumnae* (coelacanth); Q6PBL3, *Danio rerio* (zebrafish); K4GIQ5, *Callorhinchus milii* (australian ghostshark). The Figure was prepared with ESPript 3.0 (Gouet et al. 1999).

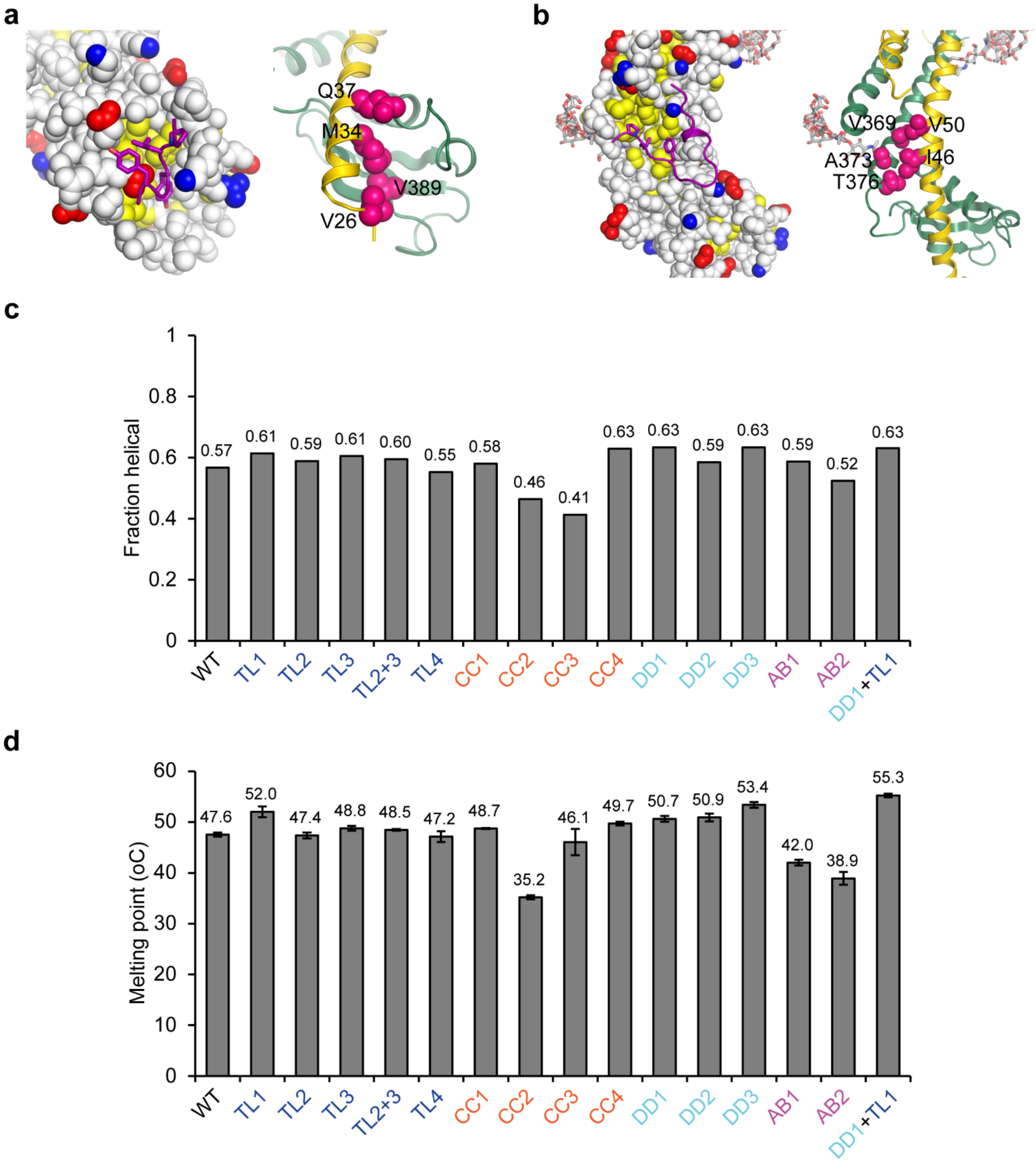

**Supplementary Figure 5: Structure-based mutagenesis of human clusterin.** (**a, b**) Mutations AB1/AB2 and CC2/CC3 of hydrophobic crystal contacts involving the α/β roll-like domain (a) and the coiled-coil (b). Contact I, mutated in AB1 and AB2 (Figure 2a), is at the α/β roll-like domain in crystal form I (a). Contact II, mutated in CC2 and CC3 (Figure 2a), is the interface of the loop containing the artificial peptide linkage (between WT-Clu residues 213 and 239) and the coiled coil domain in crystal form I (b). In each panel on the left, one Clu chain is shown in space-filling mode with positively and negatively charged residues shown in blue and red, respectively, and yellow indicating hydrophobic sidechains. Above the surface, the contacting chain segment is shown as a purple ribbon. Hydrophobic sidechains are shown in stick representation. On the right, a ribbon representation of the model is shown with the substituted residues in mutants AB1 and AB2 (a) and CC2 and CC3 (b), respectively, highlighted as pink spheres. N-glycans are shown as sticks. (**c**) Bar graph representation of estimated helical fraction of Clu mutant proteins. CD spectra were recorded at a protein concentration of 0.1 mg $ml^{-1}$ at 20 °C in 50 mM K-phosphate pH 7.0. The CD spectra from three independent experiments were averaged, molecular ellipticities calculated and the helix content (sum of H(r) and H(d)) estimated with the CONTIN algorithm. The deletion mutants TL1 and CC1, TL3 as well as most of the mutants with substitutions in or at the disulfide domain (DD1, DD3 and CC4) and the combined mutant DD1+TL1 exhibited slightly increased helix content. (**d**) Bar graph representation of melting points of Clu mutant proteins. The CD signal at 222 nm wavelength was recorded during slow heating (60 °C $h^{-1}$) of 5 µM of the respective Clu mutant protein in 50 mM K-phosphate pH 7.0. Melting points were estimated using the 'Denatured protein' function of Spectra Manager software (Jasco). Data represents averages ± SD (*n* =3 independent experiments). Most Clu variants had stability similar to WT-Clu, with melting temperatures ($T_m$) of ~47 °C. The tail deletion mutant TL1, the mutants with substitutions in or at the disulfide domain (DD1, DD2, DD3 and CC4) and the combined mutant DD1+TL1 exhibited slightly increased stability with $T_m$ values of 49.7–55.3 °C, which correlated with increased helix content (see panel c). The hydrophobic pocket mutants AB1, AB2 and CC2 showed reduced stability ($T_m$ values of 35.2–42.0 °C), suggesting that their structure was compromised by the mutations compared to WT-Clu.

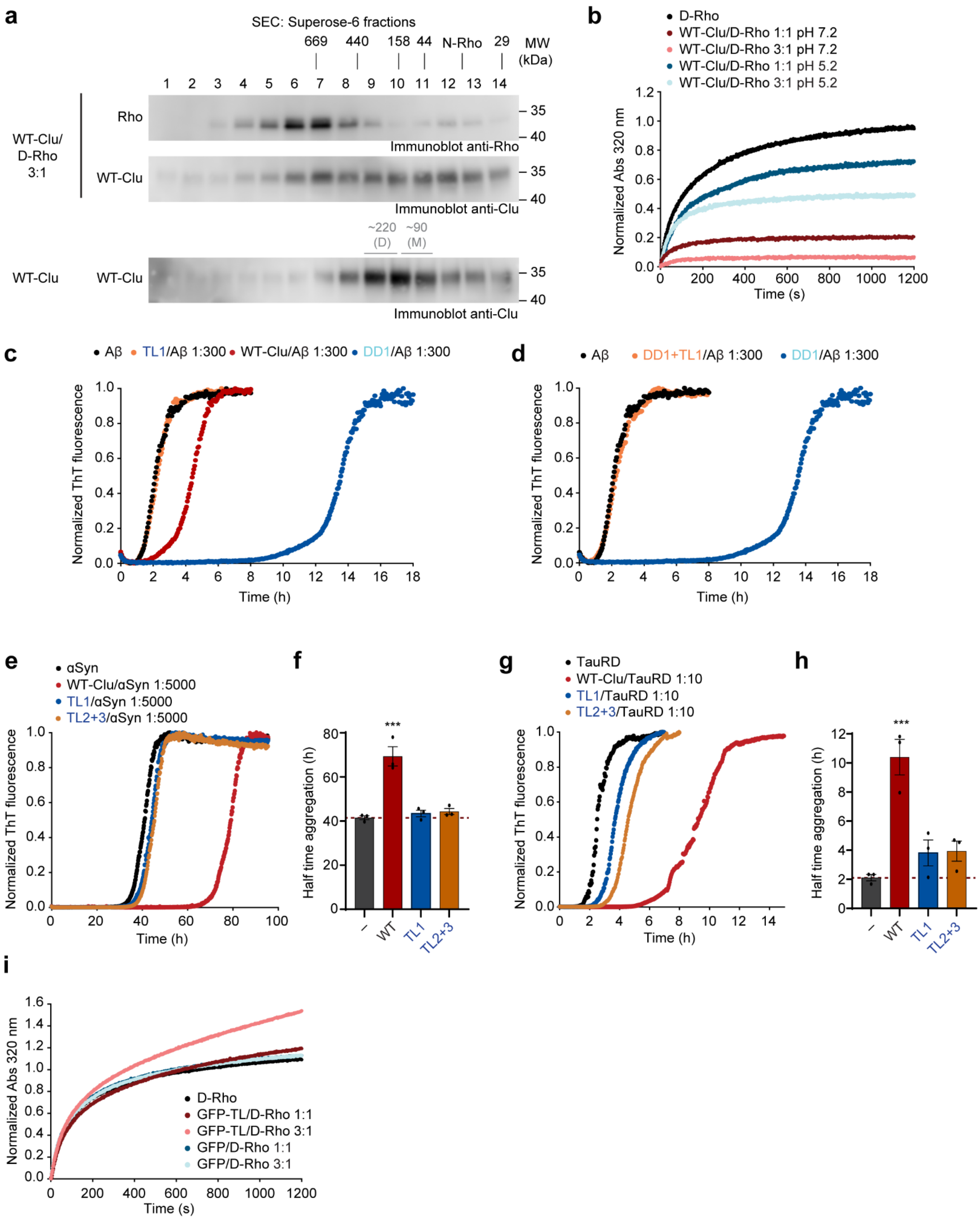
a
SEC: Superose-6 fractions
669 440 158 44 N-Rho 29 MW (kDa)
1 2 3 4 5 6 7 8 9 10 11 12 13 14
WT-Clu/ D-Rho 3:1
Rho
Immunoblot anti-Rho
WT-Clu
Immunoblot anti-Clu
~220 (D) ~90 (M)
WT-Clu
Immunoblot anti-Clu
b
D-Rho
WT-Clu/D-Rho 1:1 pH 7.2
WT-Clu/D-Rho 3:1 pH 7.2
WT-Clu/D-Rho 1:1 pH 5.2
WT-Clu/D-Rho 3:1 pH 5.2
Normalized Abs 320 nm
Time (s)
c
Aβ
TL1/Aβ 1:300
WT-Clu/Aβ 1:300
DD1/Aβ 1:300
Normalized ThT fluorescence
Time (h)
d
Aβ
DD1+TL1/Aβ 1:300
DD1/Aβ 1:300
e
αSyn
WT-Clu/αSyn 1:5000
TL1/αSyn 1:5000
TL2+3/αSyn 1:5000
f
Half time aggregation (h)
***
– WT TL1 TL2+3
g
TauRD
WT-Clu/TauRD 1:10
TL1/TauRD 1:10
TL2+3/TauRD 1:10
h
i
D-Rho
GFP-TL/D-Rho 1:1
GFP-TL/D-Rho 3:1
GFP/D-Rho 1:1
GFP/D-Rho 3:1

**Supplementary Figure 6: Clusterin substrate interactions and chaperone activity.** (**a**) Analysis of Clu–rhodanese complexes by SEC. Denatured rhodanese (D-Rho) was diluted into phosphate-buffered saline (PBS) to a final concentration of 0.5 μM in the presence of WT-Clu (1.5 μM), followed by incubation at 25 °C for 30 min. The soluble fraction obtained by centrifugation (15 min, 22,000 g) was analyzed by SEC on Superose-6, followed by SDS-PAGE and immunoblotting against Rho and Clu. A representative immunoblot is shown ($n$ =3 independent experiments). Of note, no soluble Rho remained after aggregation in absence of Clu. WT-Clu alone in PBS is shown for comparison (bottom). The retention volume of marker proteins and their molecular weight in kDa is indicated in black as well as the retention volume of native Rho (N-Rho). The apparent molecular weight of Clu alone is indicated in grey (M: monomers; D: dimers). Note that the apparent size of Clu observed by SEC is larger than expected due to its elongated shape. (**b**) Comparison of Clu holdase activity towards D-Rho at pH 7.2 and 5.2. The assay was performed as described in Figure 2b, using PBS buffer at pH 7.2 or 20 mM Na-acetate pH 5.2, 150 mM NaCl, 2 mM $CaCl_2$. Representative normalized light scattering curves of D-Rho aggregation in absence of Clu at pH 5.2 (black) or in presence of WT-Clu at pH 7.2 (Clu/D-Rho 1:1, dark red, 3:1, light red) and pH 5.2 (Clu/D-Rho 1:1, dark blue, 3:1, light blue) are shown ($n$ = 3 independent experiments). Note that maximum amplitude and kinetics of D-Rho aggregation at pH 7.2 and 5.2 were similar. (**c**) Suppression of Aβ(1–42) amyloid formation by Clu mutant DD1. Aβ amyloid formation was monitored by thioflavin-T (ThT) fluorescence in absence (black) or presence of DD1 mutant (dark blue, molar ratio Clu/Aβ: 1:300) as described in Figure 2d. Aggregation in presence of WT-Clu (red) and mutant TL1 (orange) at the same molar ratio is shown for comparison. Representative normalized fluorescence traces are shown. ($n$ = 3 independent experiments). (**d**) Effect of Clu mutant DD1+TL1 on Aβ amyloid formation. Aβ amyloid formation was monitored by ThT fluorescence in absence (black) or presence of DD1+TL1 mutant (orange, molar ratio Clu/Aβ: 1:300) as described in Figure 2d. Aβ amyloid formation in presence of mutant DD1 (dark blue) is shown for comparison. (**e-h**) Postponement of α-synuclein and TauRD amyloid formation in presence of Clu variants. Aggregation reactions containing 200 μM α-synuclein (αSyn) (e) or 10 μM TauRD (cysteine-free tau, residues 244-372 including two frontotemporal dementia mutations C291A/P301L/C322A/V337M (Yuste-Checa et al. 2021)) (g) in the absence (black) or presence of WT-Clu (red), TL1 mutant (blue) or TL2+3 (orange) (molar ratios Clu/αSyn 1:5000 and

Clu/TauRD 1:10) were monitored by ThT fluorescence. Representative fluorescence traces are shown. f, h) Bar graph showing the relative delay of αSyn (f) and TauRD (h) aggregation by WT-Clu and mutants determined by the half time of reaching the aggregation plateau (red dashed line, αSyn or TauRD alone average). Data represents averages ± SEM (*n* =3 independent experiments). *** $p<0.001$ by one-way ANOVA with Dunnett's post hoc test comparing Clu/αSyn or Clu/αTauRD to αSyn or TauRD alone. (**i**) Effect of GFP-TL on aggregation of D-Rho. The assay was performed as described in Figure 2b. Representative normalized absorbance traces of D-Rho alone (black), with additional GFP-TL (1:1, dark red; 3:1, light red) and with GFP as control (1:1, dark blue; 3:1, light blue) are shown (*n* =3 independent experiments).

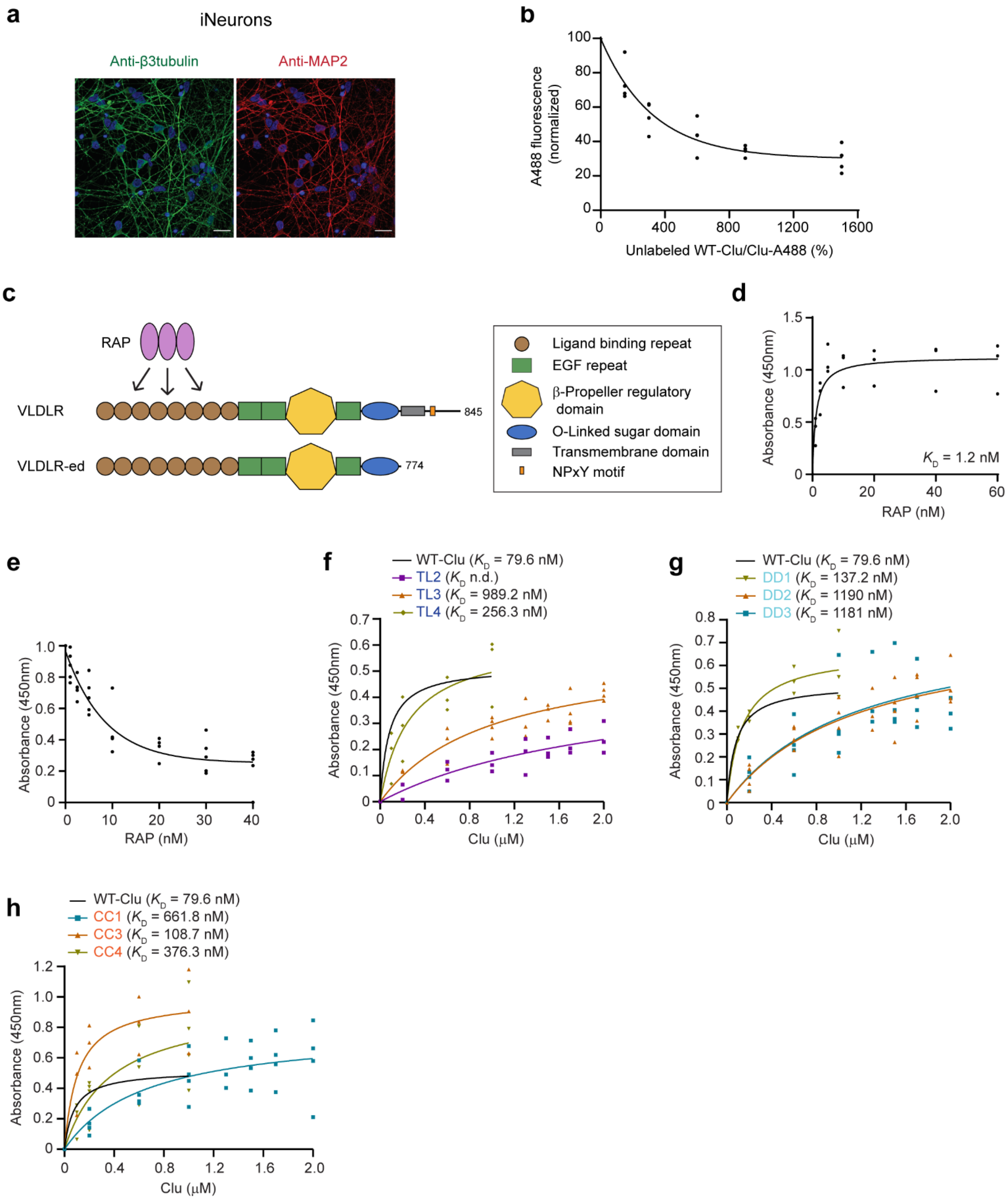

**Supplementary Figure 7: Cellular uptake and VLDLR receptor binding of WT-Clu and mutant proteins.** (**a**) Representative fluorescence microscopy of iNeurons after immunostaining for the neuronal marker proteins β3 tubulin and MAP2. The β3 tubulin and MAP2 signals are green and red, respectively. Nuclear staining with DAPI is shown in blue. Scale bars, 20 μm. (**b**) Competition of cellular uptake of Clu-A488 by unlabeled Clu. The fluorescence signal from WT-Clu-A488 internalized into iNeurons in the presence of increasing concentrations of unlabeled WT-Clu as measured by flow cytometry is shown. Competition curve is shown. Individual data points are plotted ($n = 4$ independent experiments). (**c**) Domain structure of VLDLR and the VLDLR ectodomain (VLDLR-ed) construct. The extracellular LDLR ligand binding repeats, epidermal growth factor (EGF) repeats and β-propeller regulatory domain of VLDLR are shown in brown, green and yellow, respectively. The transmembrane helix is shown in grey. The cytosolic NPxY motif (orange) serves as an internalization signal. The three repeats in low density lipoprotein receptor-related protein-associated protein 1 (RAP, magenta), which acts as a chaperone of LDLR family proteins in the ER, each recognize a pair of LDLR ligand binding repeats (Fisher et al. 2006). RAP is normally retained in the ER. (**d**) Binding curve of the interaction of RAP with VLDLR-ed as determined by enzyme-linked immunoassay (ELISA) using anti-RAP antibody. ELISA plates were coated with VLDLR-ed, and a series of RAP concentrations was added (1-60 nM). After extensive washing, bound RAP was immunodetected. RAP binding to BSA coated wells was used for a background binding correction. The $K_D$ value and binding curve are shown. Individual data points are plotted ($n = 3$ independent experiments). (**e**) Displacement of Clu from VLDLR-ed by increasing concentrations of RAP determined by ELISA using anti-Clu antibody. ELISA plates were coated with VLDLR-ed and RAP was titrated (1-40 nM) in the presence 100 nM Clu. Clu binding to BSA coated wells was considered background and subtracted. Competition curve is shown. Individual data points are plotted ($n = 5$ independent experiments). (**f-h**) Affinity of selected mutants to immobilized VLDLR-ed. Binding curves of some of the tail (f), DD mutants (g) and coiled-coil mutants (h) determined by ELISA as described in Figure 3c. The binding curve of WT-Clu is shown for comparison. The $K_D$ values and binding curves are shown (n.d., not determined). Individual data points are plotted ($n \geq 3$ independent experiments).

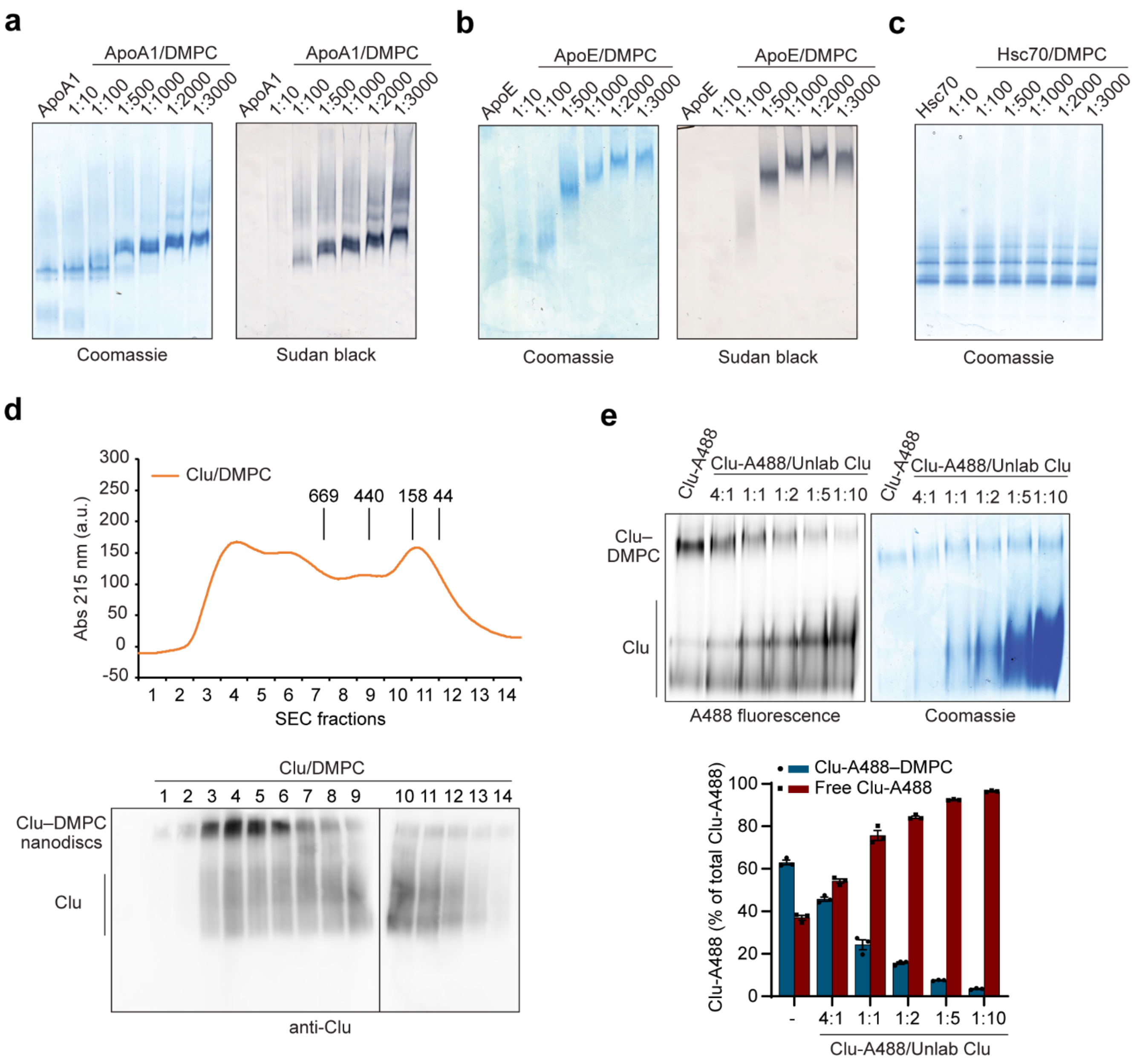

**Supplementary Figure 8: Lipoprotein particle formation and dynamics.** (**a, b**) Formation of lipoprotein particles with ApoA1 (a) and ApoE (b) detected by native PAGE. Coomassie blue (protein, left) and Sudan black (lipid, right) stained gels are shown. Apolipoprotein and DMPC, at the indicated molar ratios in PBS, were cycled above (30 °C) and below (18 °C) the melt transition temperature of DMPC (24 °C). ApoA1 and ApoE alone are shown for comparison. Representative native gels are shown ($n = 3$ independent experiments). (**c**) Incubation of Hsc70 with the phospholipid DMPC. Human Hsc70 and DMPC at the indicated molar ratios were cycled above (30 °C) and below (18 °C) the melt transition temperature of DMPC (24 °C).

Hsc70 alone is shown for comparison. A representative native gel is shown (*n* = 3 independent experiments). (**d**) Fractionation of lipoprotein particles formed from WT-Clu and DMPC by SEC. The WT-Clu/DMPC reaction mixture was analyzed by analytical SEC on a Superose-6 column equilibrated in PBS. The absorbance trace at 215 nm is shown in the top panel. Fractions were analyzed by native PAGE and immunoblotting. An immunoblot against Clu is shown in the bottom panel. Clu lipoprotein particles (Clu–DMPC) and free Clu are indicated. A representative experiment is shown (*n* = 3 independent experiments). Free Clu in fractions 3-6 was probably released from lipoprotein particles, as Clu species typically elute later (fractions 9–12). The retention volume of molecular weight markers with their respective mass (in kDa) is indicated. (**e**) Exchange of Clu from lipoprotein particles by free Clu. Purified lipoprotein particles containing Clu-A488 were incubated for 2 h in PBS at 30 °C with increasing concentrations of unlabeled Clu at the indicated molar ratios. The depletion of Clu-A488 from lipoprotein particles was analyzed by native gel and fluorescence detection (top left). A Coomassie-stained native PAGE gel is shown for comparison (top right). Representative gels are shown (*n* = 3 independent experiments). The bar graph on the bottom shows the quantification of the fluorescence signals from the Clu–DMPC lipoprotein complex (dark blue) and free Clu (dark red).

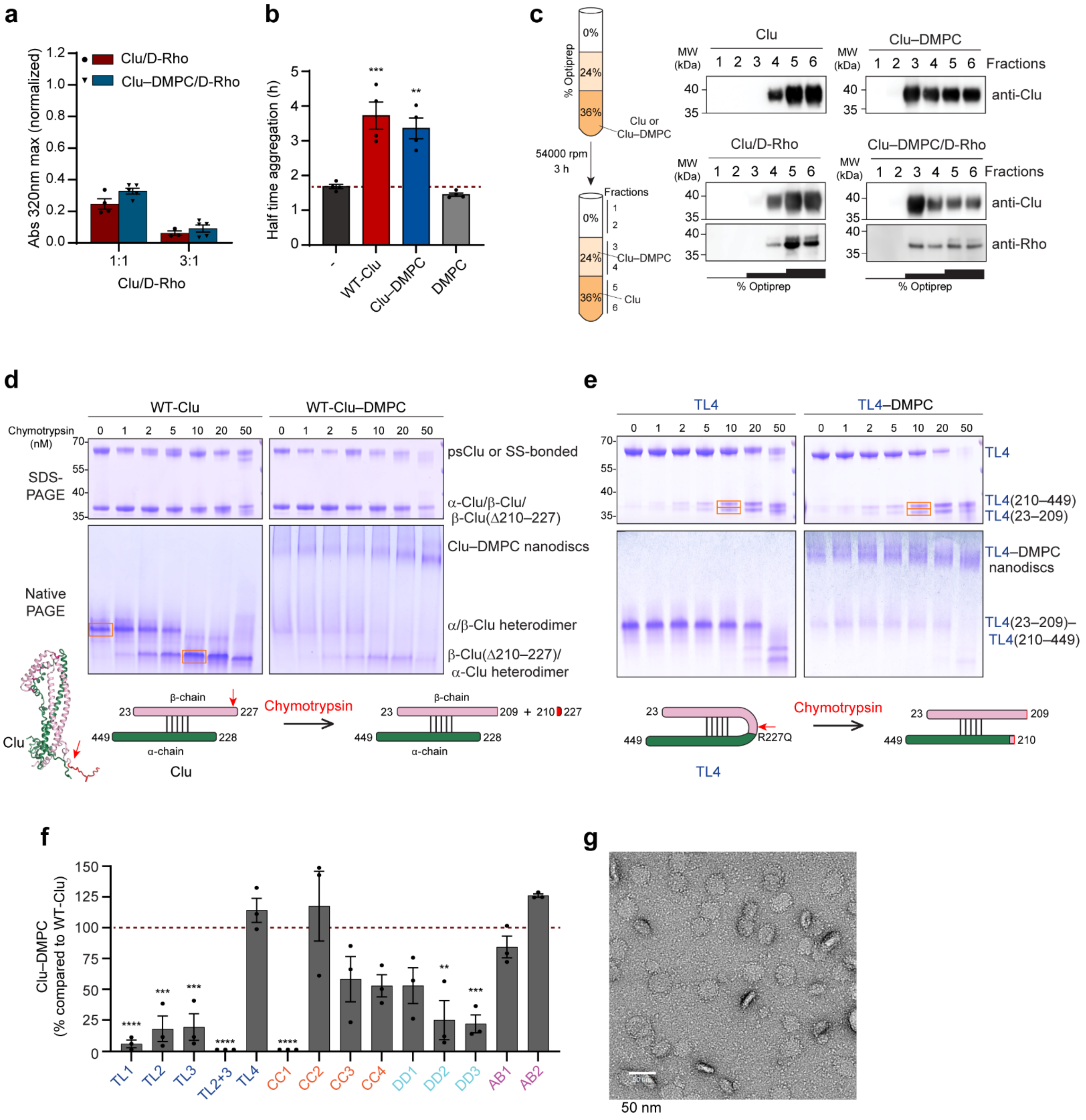

**Supplementary Figure 9: Functional and structural characterization of Clu lipoprotein particles.** (**a**) Bar graph showing the maximum plateau absorbance of D-Rho aggregation measured by turbidity assay as in Figure 2c in the presence of Clu or Clu–DMPC. Representative curves are shown in Figure 4c. Data represents averages ± SEM ($n \geq 3$ independent experiments).

(**b**) Bar graph showing quantification of Aβ amyloid formation as in Figure 2e in absence (black) or presence of WT-Clu (red), purified Clu lipoprotein complexes (Clu–DMPC, blue) or DMPC (grey) (molar ratio Clu/Aβ: 1:300 or corresponding amount of DMPC). Red dashed line, Aβ aggregation alone. Representative curves are shown in Figure 4d. DMPC alone had no effect. Data represents averages ± SEM (*n* = 4 independent experiments). ** p<0.01, *** p<0.001 by one-way ANOVA with Dunnett's post hoc test comparing Clu/Aβ to Aβ alone. (**c**) D-Rho binding to Clu–DMPC complexes by flotation assay. D-Rho aggregation reactions were performed with purified Clu lipoprotein complexes or free Clu at a molar ratio of Clu/D-Rho of 3:1 for 30 min in PBS at 25 °C. The reactions were centrifuged and the soluble fraction was mixed with Optiprep to a final concentration of 36%. 1 ml of 24% Optiprep was layered above and subsequently 1 ml of PBS on top. The density gradient stack (left panel) was ultracentrifuged and fractions of 0.5 ml were analyzed by SDS-PAGE and immunoblotting against Clu and Rho. Clu and Clu–DMPC alone were analyzed for comparison. Representative immunoblots are shown on the right (*n* = 3 independent experiments). Molecular weight (MW) standards are indicated. (**d, e**) Limited proteolysis of free protein versus lipoprotein complexes using WT-Clu (d) or the single chain mutant TL4 (e). Free protein (left) and lipoprotein particles (right) – both of the oligo-mannose form and at 10 μM Clu – were incubated for 30 min in TBS-C at 25 °C with the indicated concentrations of chymotrypsin. After inhibiting proteolysis with 2 mM PMSF on ice, the samples were analyzed by SDS-PAGE (top panels) and native PAGE (bottom panels). Coomassie-stained gels from representative experiments are shown (n=3 independent experiments). Educt and product bands are indicated. (d) While proteolytic processing of WT-Clu did not become apparent in SDS-PAGE, a large shift was visible in native PAGE for free WT-Clu. LC-MS/MS analysis of the bands indicated by red frames demonstrated that the product consists of the α-Clu/β-Clu-Δ(210-227) heterodimer linked by five disulfide bonds (see scheme below). The most sensitive peptide bond (209-210) is located in the β-tail (see structure representation). The large shift in native PAGE is apparently caused by a change in net charge due to loss of positively charged residues in the β-tail. Of note, the peptide Clu(210-227) (not resolved) bears one Lys and four Arg residues and no Asp/Glu. (e) Limited proteolysis of both free TL4 (left) and TL4–DMPC (right) results in two bands near 35 kDa in the SDS-PAGE gels. Proteomics analysis of the bands indicated by red boxes demonstrated that they corresponded to the fragments TL4(23-209) and TL4(210-449), i.e. the same site is most sensitive to chymotrypsin in both WT-Clu and TL4. Because proteolytic nicking (see scheme) does not change the net charge, the electrophoretic mobility of the cleaved product band is identical to that of TL4 in native PAGE. (**f**) Analysis of lipoprotein complex formation by Clu mutants. Formation of lipoprotein complexes with Clu mutant proteins was monitored by native PAGE at 1:500 molar ratio (Clu–DMPC), and the Coomassie blue-stained bands quantified by densitometry (representative gels are shown in Figure 4e). The bar graph shows the relative intensity of the mutant lipoprotein complex band compared to WT-Clu (dashed line). Data represents averages ± SEM (*n* =3 independent experiments). ** p<0.01, *** p<0.001 and **** p<0.0001 by one-way ANOVA with Dunnett's post hoc test comparing Clu mutants to WT-Clu. (**g**) Analysis of lipoprotein complexes formed by the GFP fusion protein αTL-H7 by negative stain electron microscopy. A representative micrograph of a αTL-H7 lipoprotein complex preparation performed as in Figure 4f is shown (*n* =3 independent samples). The nanodiscs are roughly 25–45 nm in size. The nanodisc rims are decorated with tiny dots, presumably the GFP moieties. Scale bar, 50 nm.

**Supplementary Table 1: Crystallographic data collection and refinement statistics.**

| | Crystal form I | Crystal form II** |
|---|---|---|
| **Data collection** | | |
| Space group | $P2_1$ | $C2$ |
| Cell dimensions | | |
| $a$, $b$, $c$ (Å) | 65.74, 43.81, 102.80 | 194.44, 46.44, 155.17 |
| $\alpha$, $\beta$, $\gamma$ (°) | 90, 107.29, 90 | 90, 127.20, 90 |
| Resolution (Å) | 2.8 (2.95 – 2.8) * | 3.5 (3.83 – 3.5) * |
| $R_{merge}$ | 0.127 (1.360) | 0.200 (1.413) |
| $I$ / $\sigma I$ | 5.9 (0.7) | 6.1 (1.4) |
| Completeness (%) | 98.2 (96.1) | 99.8 (99.6) |
| Redundancy | 3.7 (3.6) | 7.2 (5.7) |
| **Refinement** | | |
| Resolution (Å) | 2.8 | 3.5 |
| No. reflections | 13814 | 14331 |
| $R_{work}$ / $R_{free}$ | 0.2322 / 0.2742 | 0.2352 / 0.2836 |
| No. atoms | | |
| Protein | 3007 | 5726 |
| Ligand/ion | 123 | 238 |
| Water | – | |
| $B$-factors | | |
| Protein | 79.42 | 135.42 |
| Ligand/ion | 119.48 | 182.16 |
| Water | – | – |
| R.m.s. deviations | | |
| Bond lengths (Å) | 0.003 | 0.003 |
| Bond angles (°) | 0.522 | 0.505 |

**Data from two crystals were merged for processing of crystal form II. *Values in parentheses are for highest-resolution shell.

**Supplementary Information References**

Choi-Miura, N. H., et al. (1992). "Identification of the disulfide bonds in human plasma protein SP-40,40 (apolipoprotein-J)." J Biochem **112**(4): 557-561.
Fisher, C., et al. (2006). "Structure of an LDLR-RAP complex reveals a general mode for ligand recognition by lipoprotein receptors." Mol Cell **22**(2): 277-283.
Gouet, P., et al. (1999). "ESPript: multiple sequence alignments in PostScript." Bioinformatics **15**: 305-308.
Yuste-Checa, P., et al. (2021). "The extracellular chaperone Clusterin enhances Tau aggregate seeding in a cellular model." Nat Commun **12**(1): 4863.
